# Rapid Synthesis of Cryo-ET Data for Training Deep Learning Models

**DOI:** 10.1101/2023.04.28.538636

**Authors:** Carson Purnell, Jessica Heebner, Michael T. Swulius, Ryan Hylton, Seth Kabonick, Michael Grillo, Sergei Grigoryev, Fred Heberle, M. Neal Waxham, Matthew T. Swulius

## Abstract

Deep learning excels at cryo-tomographic image restoration and segmentation tasks but is hindered by a lack of training data. Here we introduce cryo-TomoSim (CTS), a MATLAB-based software package that builds coarse-grained models of macromolecular complexes embedded in vitreous ice and then simulates transmitted electron tilt series for tomographic reconstruction. We then demonstrate the effectiveness of these simulated datasets in training different deep learning models for use on real cryotomographic reconstructions. Computer-generated ground truth datasets provide the means for training models with voxel-level precision, allowing for unprecedented denoising and precise molecular segmentation of datasets. By modeling phenomena such as a three-dimensional contrast transfer function, probabilistic detection events, and radiation-induced damage, the simulated cryo-electron tomograms can cover a large range of imaging content and conditions to optimize training sets. When paired with small amounts of training data from real tomograms, networks become incredibly accurate at segmenting *in situ* macromolecular assemblies across a wide range of biological contexts.

**Summary:** By pairing rapidly synthesized Cryo-ET data with computed ground truths, deep learning models can be trained to accurately restore and segment real tomograms of biological structures both *in vitro* and *in situ*.

## Introduction

Cryo-electron tomography (cryo-ET) allows the direct three-dimensional imaging of purified macromolecules, enriched organelles, whole bacterial and archaeal cells, and eukaryotic cellular compartments in a frozen-hydrated state ^1–11^. It is only a matter of time before all biological material is subjected to its investigation ^12–16^. Cryotomograms can be rich with information across length scales spanning a few angstroms to several microns ^17,18^, but extracting all of this information automatically (if at all) is challenging given the variation in their content and complexity.

Deep learning has emerged as a powerful tool for image restoration ^19–21^ and segmentation ^22–25^ in cryo-ET, but perfect ground truth datasets do not exist and cannot be generated experimentally. Collecting training data at the microscope is both expensive and slow, as well as a drain on valuable time that could be spent collecting new experimental datasets. Another obstacle is that a single tomogram rarely contains enough views from each feature-class to train a reliable model for their detection. Data augmentation is an established approach to increasing the dataset depth of variability (^26,27^), but the missing-wedge generates orientation-specific distortions in tomographic data, complicating this sampling problem. Finally, even if one has a good set of tomographic data to train on with adequate signal and a full angular sampling, it will need manual annotation into a segmentation for supervised learning.

Simulating accurate training data and inferring these networks to real data is an appealing solution ^28^. With simulated data, users could quickly generate new mixtures of molecules from existing structural models, explore a range of imaging parameters, and produce perfect ground truth segmentations all in a matter of minutes. Training with simulated cryo-ET data is a novel approach to overcoming limitations with deep learning, and currently, no software is dedicated to this objective. To this end, we have developed cryo-TomoSim (CTS), a MATLAB-based program that simulates tilt series from coarse-grained models of molecular mixtures in vitreous ice for tomographic reconstruction. CTS provides easy control of a variety of modeling and simulation parameters that allow users to quickly generate a wide range of training data from any number of input structures. We then go on to demonstrate the effectiveness of CTS-generated datasets in training both regression denoising and semantic segmentation U-Net models.

## CTS Overview

### Software Environment

We developed CTS in MATLAB, because it is a well-documented platform that is available on all mainstream operating systems. Matlab code is usually transparent and readable, so tools using it are relatively easy to modify for personal use cases. CTS can be controlled from either the MATLAB command line or through a dedicated graphical interface, making it accessible to nearly anyone with a modern computer. It calls on the well-known software suite IMOD ^29^ to rapidly generate projection images and automatically reconstruct simulated tilt series into tomograms, and then organizes them into directories containing data from multiple stages of the simulation process. Finally, CTS generates a ground truth atlas at the dimensions of the tomogram, with the voxels belonging to each feature-class precisely annotated.

### Modeling schema

#### Coarse-grained Model Generation

CTS generates models as voxel-based constructs based on a specified pixel size and 3D volume, rather than generating a full atomic map. This permits low computational times and the ability to run on any modern computer, without requiring GPU acceleration, many CPUs, or large amounts of RAM. The majority of development and testing was done on a low-spec laptop running on an intel i5 CPU with 4 cores and 8GB ram, which takes 5-10 minutes to run CTS for typical models from structure inputs to a complete reconstruction.

Individual structure files (pdb or cif, Fig. 1A) are converted into coulomb potential maps, at the specified resolution, through the simple accumulation of atomic potentials in each voxel, approximated as the atomic number of the atom. As the model does not retain any atomic information and the simulation is performed at the voxel level, dispersion is not modeled due to its negligible impact at the resolution of a single cryo-electron tomogram.

**Figure 1.**
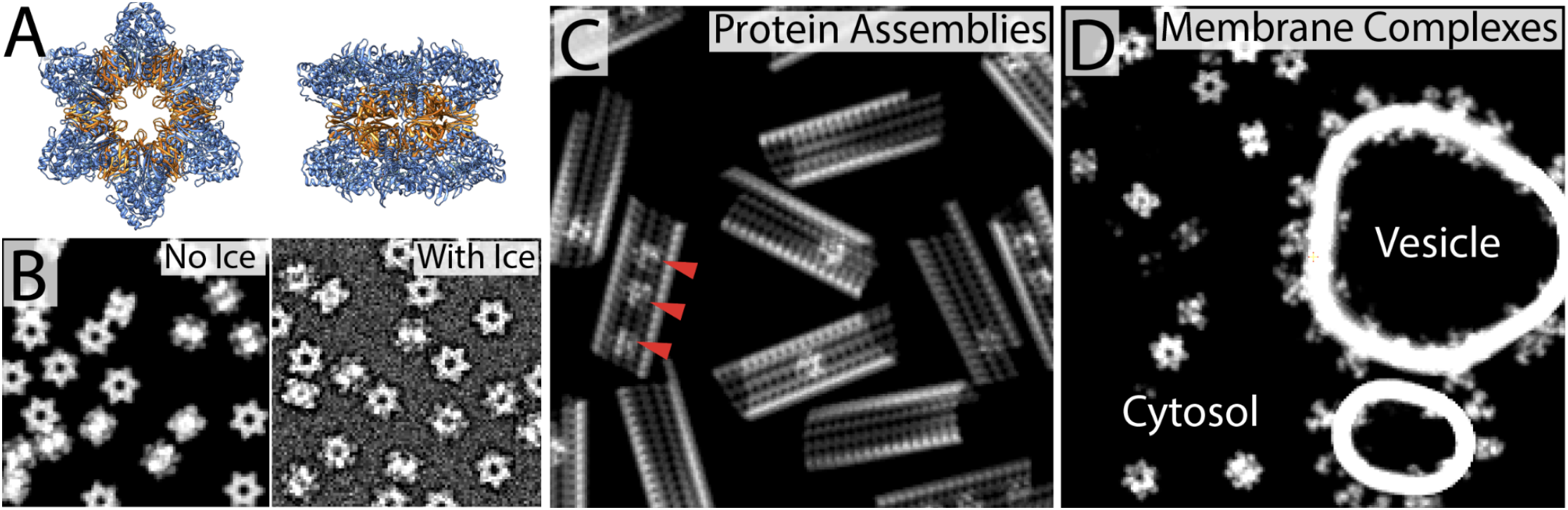
Modeling with CTS A) Two views from the atomic model of the dodecameric alpha-CaMKII holoenzyme, colorized by its catalytic (blue) and associative (orange) domains. B) Voxel based models of fields of CaMKII without and with ice embedding. C) CTS model built with microtubule “assembly” containing three intralumenal particles (red arrow heads). During placement of each microtubule, the number of intralumenal particles was randomly determined. D) CTS model of membranous vesicle decorated with NMDA receptors and a GPCR. CaMKII was flagged as a cytosolic protein before modeling.

#### Handling of different particle schemes

CTS modeling can be as simple as inputting any arbitrary number of independent structure files, but has support for several types of structured arrangements or special handling. For instance, clusters of unconstrained proteins, as well as bundles of linearly oriented proteins, can be generated to create different orders of particle packing (Fig. S1). Protein complexes can also be modeled, including partial occupancy of secondary members on a primary scaffold (Fig. 1C). Complexes also extend to subcomponents of an individual protein. Particles can be placed in association with, including transmembrane passage through, lipid membranes. They can also be restricted exclusively to inside or outside of vesicles (Fig. 1D).

#### Model constraints

After choosing input particles, model constraints must be set, such as the volume dimensions and features. If desired, constraints can be placed on the volume to determine which outside borders, if any, particles can cross during model filling. The default being the top and bottom of the Z plane to mimic cryo preparation flattening a sample within a thin layer of vitreous ice, while allowing proteins to clip out of the model area in other directions.

Optionally, CTS models can include portions of a carbon support as well as vesicles (Fig. 2A) prior to model filling. Both are modeled similarly to proteins from structure files in that they are generated by accumulation of density from particles within each voxel. The carbon support is a perfectly flat plane of amorphous atomic carbon with a circular hole, offset so that one edge of the model is occupied by carbon (Fig. 2A). Vesicles are modeled as lipid density corresponding to a bimodal distribution across the radius of the vesicle shape, which can range from perfectly spherical to highly irregular. After model filling, CTS can also optionally place spherical gold fiducials (Fig. 2A) into the model before adding vitreous ice. Ice is generated as a global field of vitreous water applied as minimal values to voxels (Fig. 1A), while fiducials are gold atoms randomly generated in a sphere. As it would be computationally prohibitive, individual atoms are not randomly generated within these distributions but instead pseudo-atoms that contribute proportionally higher density at lower model resolution.

**Figure 2.**
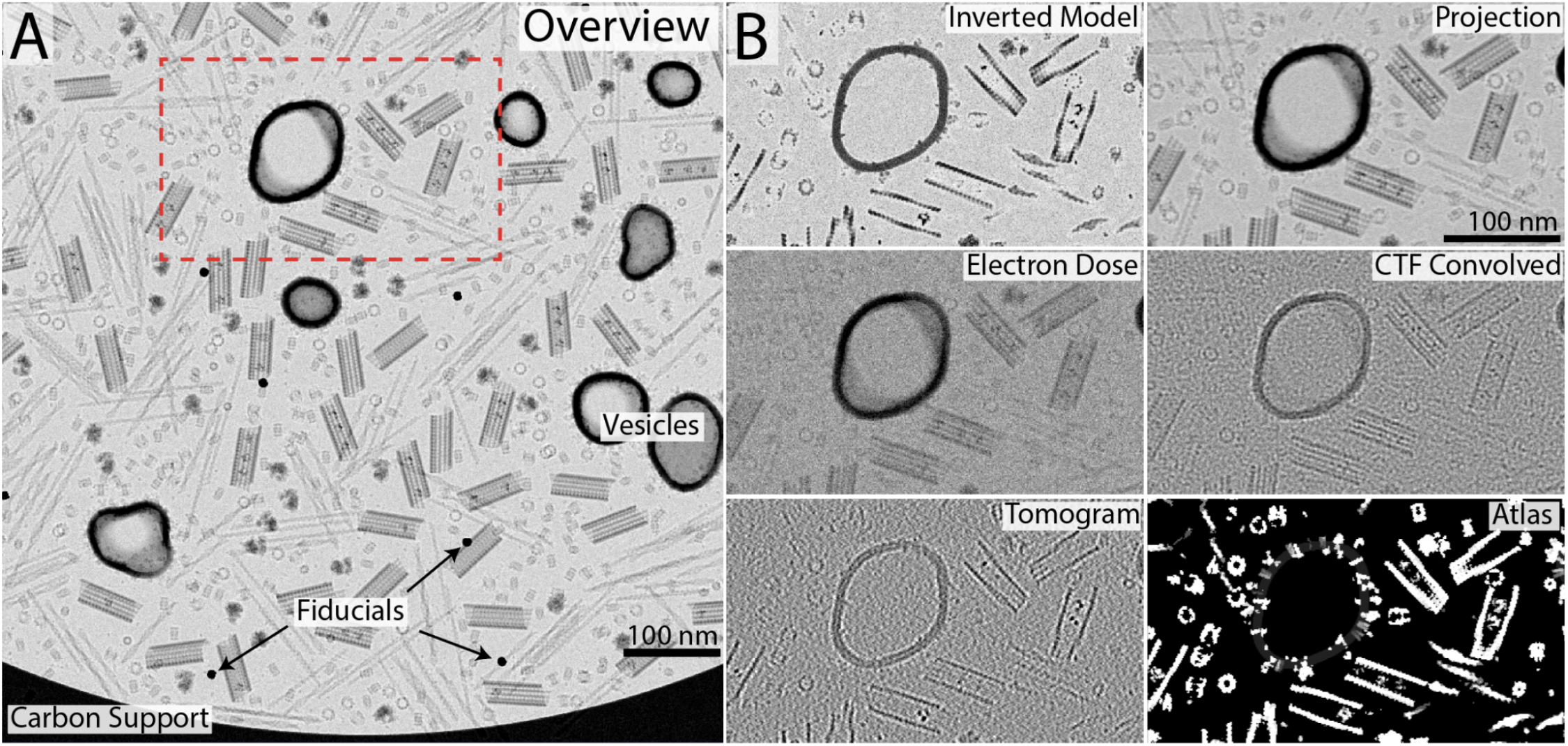
Simulating data with CTS. A) Simulated projection through tomographic model. B) Detailed stages of output from the CTS workflow focused on the boxed region in (A). The progression is Model, Projection, Electron Dose, Contrast Transfer Function (CTF), and then Tomogram. The ground truth atlas is the final component generated.

#### Brute force model filling

After constraints and generation of initial features, the model is filled with the selected input particles (Fig. 1A). Particles are organized into several different ‘layers’ that are used for model filling in succession, to allow users more precise control over particle mixtures when desired. Filling is conducted in a brute-force approach, with placements attempted iteratively at semi-random locations not known to be occupied. Input particles are randomly selected, rotated, and placed into these coordinates. If this causes an overlap with an existing object, it is rejected and the process continues until a maximum density or number of iterations is reached. Early layers therefore occupy more of the volume than successive layers, the last of which is often useful for inserting small ‘distractor’ particles without crowding out larger structures of interest. After model filling, fiducials and ice are added as described above. It is important to note that in addition to the final combined model, CTS generates and stores per-class models of what is placed into the combined model that is later used to generate an atlas of all simulation contents.

### Simulation schema

Proceeding from a complete model, the simulation is also conducted on a per-pixel level for speed, and so does not simulate image formation at the precision of an electron wave interacting with and propagating through individual atoms.

#### Inputs

The CTS simulator has many parameters, covering all reasonable and many unreasonable as well as technically impossible imaging capabilities. The control parameters are the same as standard imaging parameters: tilt increment and limits, defocus, electron dose, and tilt scheme (Fig. S2 & S3). Advanced options include control over radiation damage, deviation from exact tilt angles, inelastic electron scattering, and generation of “ideal” images that lack CTF and dose sampling.

#### Projecting tilt series with IMOD

The first step of simulation is inversion of the input model contrast to match the standard dark-on-bright density scale generated by cryo-EM (Fig. 2B, model). The inverted model is then projected as a tilt series using the *xyzproj* command from the IMOD ^29^ package along the specified tilt axis, at the specified tilt angles (Fig. 2B, Projection). CTS defaults expect balanced tomographic tilt series but can project any arbitrary set of tilts in any order, and so can also generate single micrographs.

#### Dose sampling

Image generation is simulated by sampling electrons from a distribution of the tilt angle’s scattering potential (Fig. 2B, Electron Dose). For each tilt, the camera-detected dose is adjusted based on the maximal DQE of the detector as well as inelastic scattering of electrons away from the path of the detector. This dose-adjusted scattering map is used as the lambda parameter of a poisson distribution from which detected electrons are drawn. The following equation is used to determine transmitted dose:

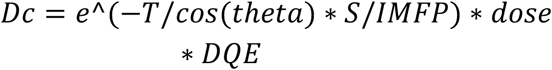

*Dc* is the corrected dose transmitted, *S* is the scattering factor (1), *IMFP* is the inelastic mean free path of vitreous ice (3.8nm), *T* is the thickness of the sample, *theta* is the tilt angle, and *DQE* is the detector quantum efficiency of the camera.

#### CTF intensity modulation

Once tilt images are projected, they are modulated by a contrast transfer function (CTF) ^30^ based on the pixel size and defocus, as well as microscope parameters such as the accelerating voltage and spherical aberration (Fig. 2B, CTF). The CTF in Fourier space is computed according to the following functions:

1. *CTF* = *E* ∗ ((1 − *Q*)*sin*(*eq*) + *Qscos*(*eq*))
2. *eq* = *pi*/2 ∗ (*CS* ∗ *L*^3^ ∗ *k*^4^ − 2 ∗ *Dz* ∗ *L* ∗ *k*^2^)
3. 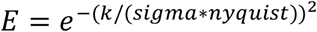
4. *L* = *H* ∗ *c*/*sqrt*(*e* ∗ *V* ∗ (2 ∗ *m* ∗ *c*^2^ + *e* ∗ *V*))

Overall equation for the CTF profile (1), wave component equation (2), envelope function of the contrast (3), and the calculation of the relativistic electron wavelength (4).

Where L is the relativistic electron wavelength, CS is spherical aberration, K is the spatial frequency, Dz is defocus, sigma is the envelope factor (.9), and Q is the amplitude contrast factor (.07). In 4, H is the planck constant, e the electron charge, c the speed of light, m the electron mass, and V the acceleration voltage.

To account for variable defocus across a tilted sample, CTF modulation is applied to each tilt angle in a series of overlapping strips with an adjusted defocus value. After inversion from fourier space, the strips are combined by weighted average to form a seamless composite.

#### Radiation damage

Radiation damage is modeled in a very simplified fashion. Before electrons are sampled for each tilt angle, that tilt projection is corrupted by two operations scaled by the cumulative electron dose transmitted. The first operation is a smoothing step that reduces signal clarity in higher-resolution images, and the second is a layer of gaussian noise applied to the whole tilt image.

#### Tomographic reconstruction

As CTS executes a perfectly aligned simulation with complete information, reconstruction of the tomogram requires no prior alignment steps or preprocessing. CTS performs reconstruction of the simulated tilt series (Fig. 2B, Tomogram) using IMOD’s *tilt* command and supplying the exact tilt angles, followed by using the IMOD command *trimvol* to rotate the tomographic reconstruction to a standard orientation. At present, CTS does not include any CTF-correction step for the output tomogram.

## Results

### Regression-Based Tomogram Restoration

Cryo-ET image restoration by deep learning has been the focus of several studies in the past few years ^19–21^. These approaches have found sophisticated ways to denoise without regression methods ^31^, because a true image “prior” cannot be experimentally derived in cryo-EM, due to the destructive nature of the beam. This is not true of CTS, which can generate multiple tomographic outputs from the same model, including a noise-free prior collected on a perfect “detector”.

To make a generalizable cryo-ET regression network, we used CTS to produce three small tomograms of synthetic cytoplasm (600 × 600 × 60 voxels each at 13.2 ang/pix, Fig. S4A), which took approximately 10 min each. This “cytoplasm” included a mixture of large and small macromolecules commonly found in cells as well as simulated lipid vesicles, all tightly packed like that seen in real cellular tomograms (See Table S1, S2 and S3 for all structures, modeling parameters, and simulation parameters used). To train the regression network, ideal tomogram “priors” (Fig. S4B) were paired with noisy synthetic tomograms from the same model as inputs (Fig. S4C), and network performance was validated by regressing the synthetic tomogram back to the prior state (Fig. S4D). The same network was then used to radically denoise a variety of real cryotomograms, including cross-linked actin-filaments *in vitro*, rat neuronal cytoplasm, and whole bacterial cells (Fig 3 A-C). When compared with established methods (nonlinear anisotropic diffusion and noise2noise deep learning), it is clear that regression training against CTS priors is superior in terms of denoising while also retaining structural detail (Fig. S5).

**Figure 3.**
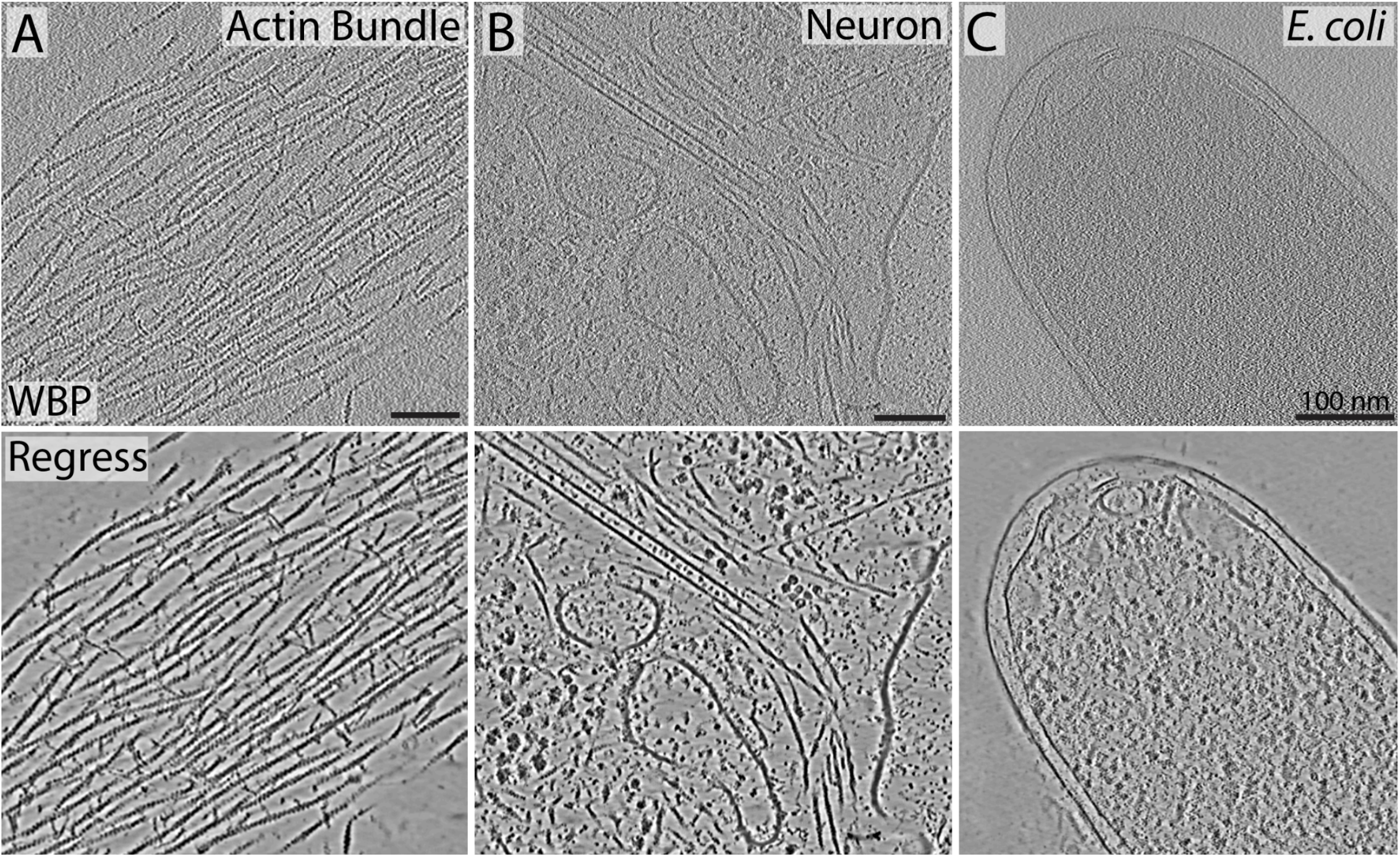
General effectiveness of CTS-trained regression networks. A-C) Deep learning-based regression by the same network on actin filaments bundled by the cross-linking protein alpha-actinin *in vitro*, Neuronal cytoplasm, and an *E. coli* cell, respectively.

Additionally, more constrained training sets can be generated quickly and easily for any *in vitro* purified sample. For instance, by simulating tomograms from a mixture of nucleosome and chromatin models (see Methods) we were able to restore tomograms of purified human chromatin to a nearly noiseless state, as well as restore the large tilt increments (5°) and missing wedge in Fourier space (Fig 4A & B).

**Figure 4.**
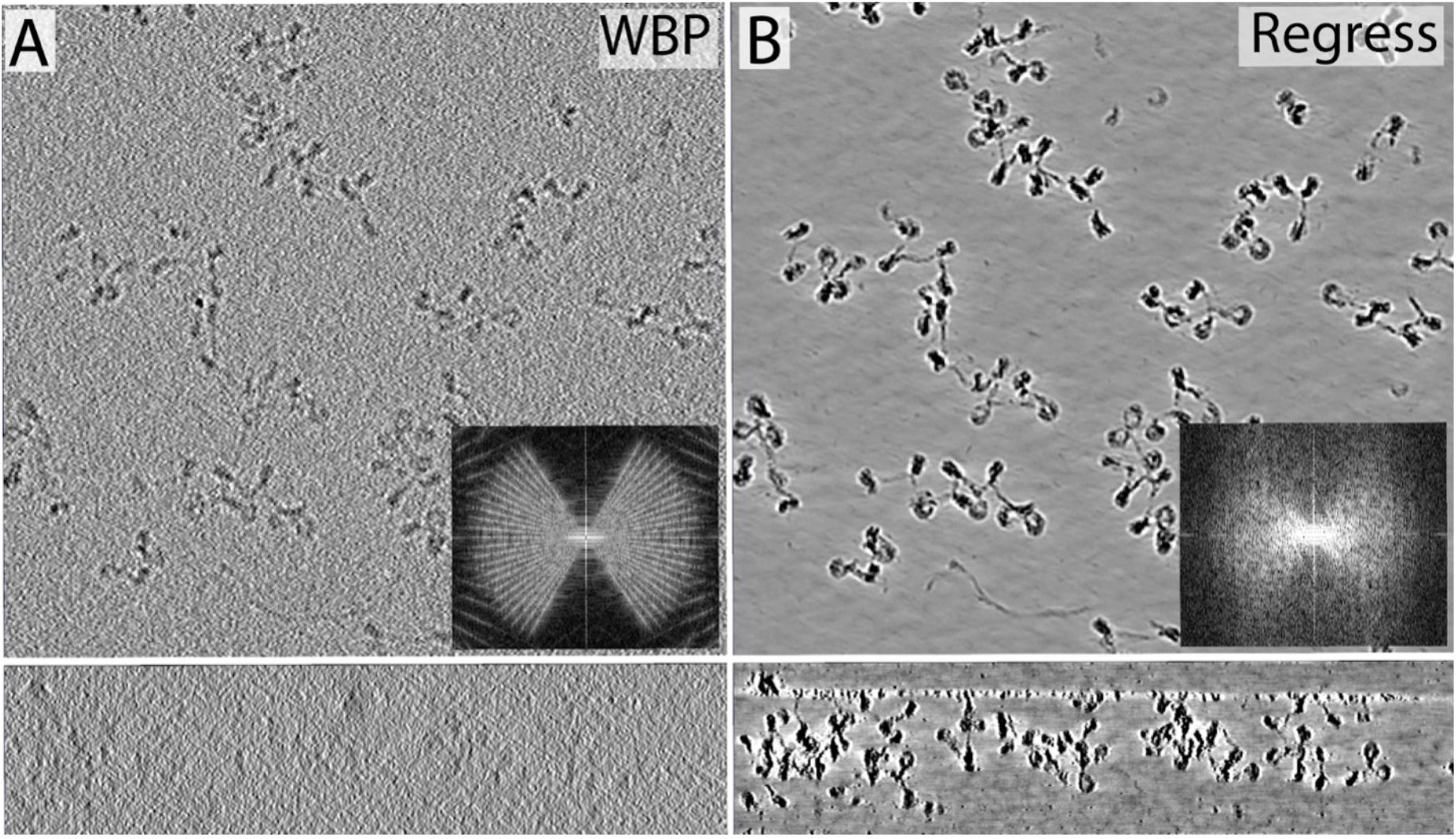
CTS-based regression on real tomograms of purified chromatin A) 100-voxel (84 nm) thick slab from a weighted back projection (WBP) cryotomogram of purified human chromatin viewed in the XY (top panel) and XZ plane (bottom panel). B) Identically sized slab from the same tomogram after regression denoising. The insets in (A) and (B) are Fourier transforms of the images in their respective lower panels.

Structural features in regression restorations are subject to bias from the U-Net transformation ^32^, so we compared the outputs from differently trained networks on the same synthetic tomogram generated from an elongated F-actin model (Fig. S6). When compared to the ground truth of the simulated weighted back projection input, all of the trained networks denoised the actin filaments well while maintaining the lower resolution details about filament helicity and actin monomer localization. It is obvious, however, from the network that has seen no actin filaments, that some of the information about monomer-to-monomer interactions is lost, as might be expected.

### Semantic Segmentation of Cryo-ET Data

Segmentation of cryo-ET data into known components is a laborious task, and even if you employ deep learning to help in the process it still requires careful manual annotation of the training data as inputs. With CTS the annotated ground truth atlas only takes seconds to produce and can be used as input for training segmentation U-Nets on the corresponding synthetic tomograms. Given that CTS can quickly derive new datasets under different “imaging” conditions, we took this opportunity to test the effects of key parameters of tomographic data collection on the inference of our trained U-Nets during segmentation.

We tested the effects of moderate changes in defocus (*μm), tota*l electron dose (−e/Å^2^), pixel size (Å/pix), and tilt increment (°) on segmenting simulated tomograms of either actin and cofilactin filaments or CaMKII divided into submodels containing the catalytic and association domains ^33^. We also quantified the effects of “missing wedge” orientation with respect to the filament’s long axis on segmentation accuracy. All networks were trained with an equal amount of data (including data augmentation parameters), for the same number of epochs to ensure fair evaluation. All segmentations were then compared to the CTS-generated ground truth and a Dice coefficient was calculated to evaluate network success. We found, under our training conditions, that U-nets were robust across all the tested parameters (Figs. S7-S10), except for the orientation of the missing wedge (Fig. S11). This makes sense given that actin filaments oriented parallel to the tilt-axis look dramatically different compared to those aligned perpendicularly. Because of this, networks trained only on datasets with filaments in one orientation with respect to the missing wedge largely fail when inferred to any dataset that contains filaments in another orientation (Fig. S11). Therefore, it is important that training data contains structures with a broad angular distribution with respect to the missing wedge for robust, accurate inference, which is exactly what CTS was designed to provide.

### Segmenting *In Vitro* Data

To segment cryotomograms of purified molecules captured in a thin sheet of vitreous ice we found that training with simulated data alone was surprisingly powerful. In an initial validation test, cryo-ET data was collected on purified actin filaments mixed with the actin binding protein cofilin, which formed a mixture of F-actin and cofilactin, a hyper-twisted filament formed by a 1:1 stoichiometric binding of cofilin to F-actin ^34^. In real cryotomograms, stretches of bare actin were clearly distinguishable from stretches of cofilactin (Fig. 5), because their helical repeat lengths are quite different (∼37 nm compared to ∼27 nm, respectively ^35^).

**Figure 5.**
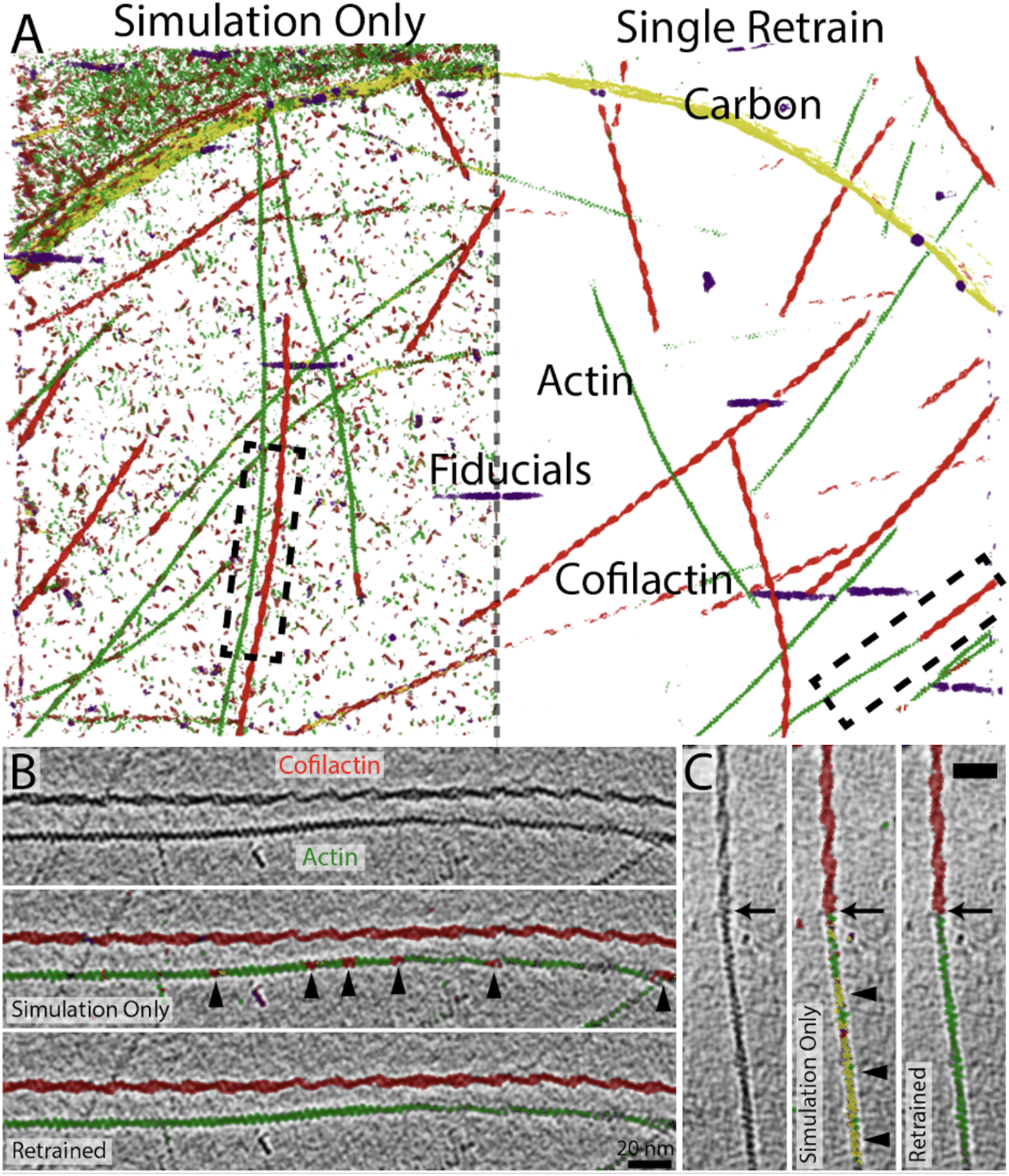
Segmentation of real *in vitro* tomograms by synthetically trained U-Nets. A) Full 3D Segmentation of a real cryotomogram containing a mixture of actin (green) and cofilactin (red) filaments. The left side shows the results from a network trained only on CTS-derived tomograms containing actin, cofilactin, fiducials (purple) and a carbon support (yellow), and the right side shows segmentation results after a single retraining with 5 hand-corrected slices. B) and C) Higher magnification views of the real tomographic data and its segmentations by both networks. The segments shown correspond to the areas surrounded by dashed boxes in (A).

To start, a 5-slice U-Net was trained against a 400 × 400 × 50 voxel CTS-generated cryotomogram containing actin filaments, cofilactin, and 10-nm gold fiducials. The edge of a synthetic carbon support was also present because the real data contained this feature. Filaments were constrained within the 50-voxel z-height to ensure they lie mostly flat, as they are found within the real tomograms. Figure 5A depicts the segmentation of the entire tomogram by two related 4-class networks. One trained by only simulated data (“simulation only”), and one further trained on 5 hand-corrected slices of the real tomogram. By examining the “simulation only” segmentation up close, it is clear that the majority of misclassified densities are small but real protein densities associated with the air-water interface, and they are almost entirely eliminated from the background in the retrained network. There were small and uncommon misclassifications along filaments (Fig. 5B & C), but these were also eliminated with a single retrain of the network. Interestingly, while there were no transitions from actin to cofilactin within individual filaments in our training set, in the real data these transitions were abundant and easily detected by the network (Fig. 5C).

To test whether this approach is generalizable and to push its limitations, we used purely synthetic datasets to not only distinguish between actin and cofilactin, but to also segment the cofilin molecules decorating the filament (Fig. 6A). We then used a similar approach to 1) differentiate between the catalytic domains and the associative core domains in real tomograms of purified alpha-CaMKII (Fig. 6B), 2) segment histone proteins from the DNA strand in purified chromatin (Fig. 6C), and 3) target the outer alpha-rings and inner beta-rings of the 20S proteasome (Fig. 6D, EMDB 7152).

**Figure 6.**
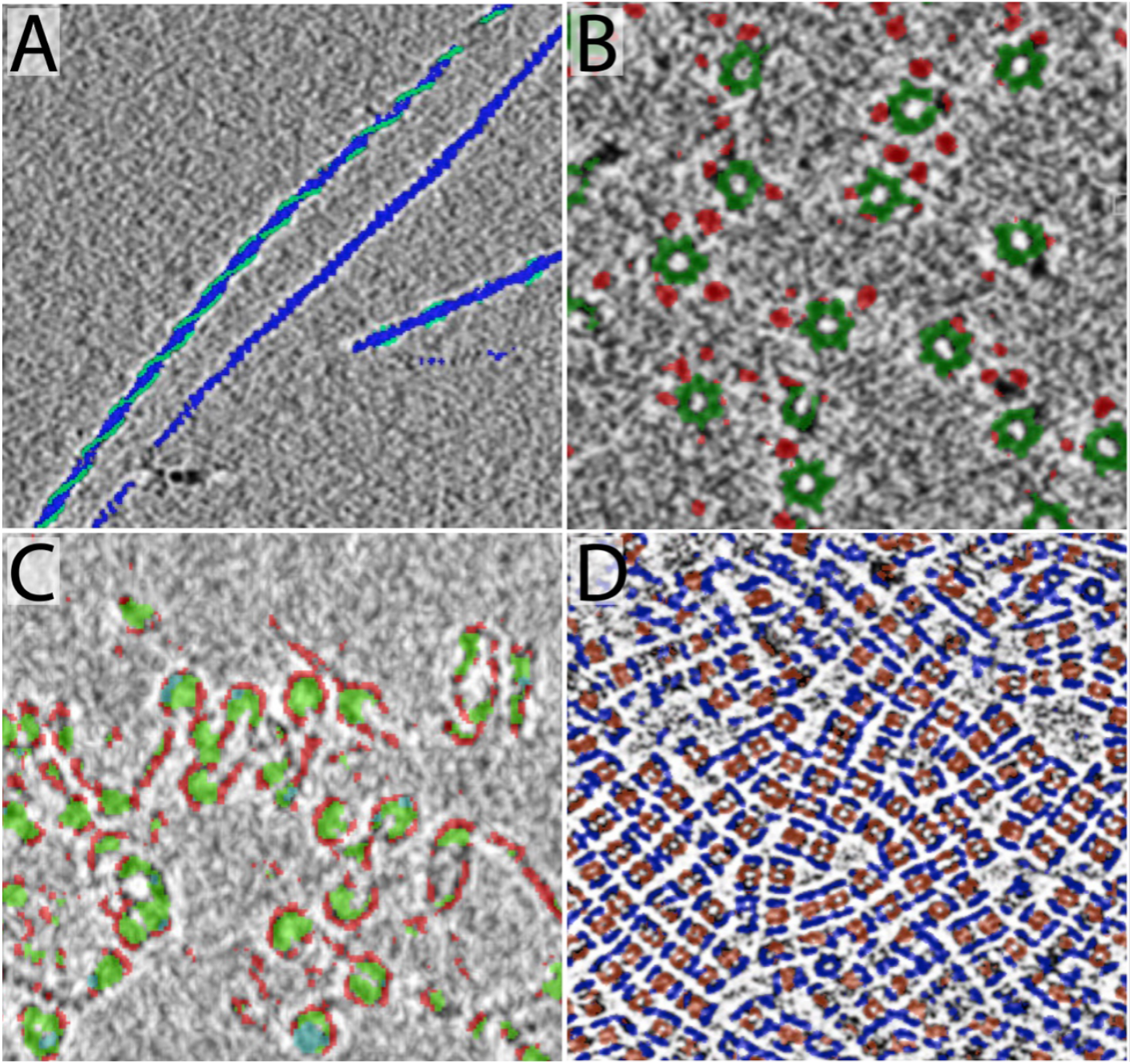
Submodel discrimination with CTS-generated training data. A) segmentation of actin subunits (blue) and cofilin subunits (teal) in a mixed in-vitro population of bare actin and cofilin-decorated actin. B) segmentation of CaMK2 alpha into central association domains (green) and variously extended catalytic domains (red). C) segmentation of low-compaction chromatin, DNA in red and histone cores in green. D) segmentation of proteasomes into terminal alpha rings (blue) and middle beta rings (orange).

### Segmenting Cellular Data

While there are many applications for segmenting *in vitro* systems, we ultimately need tools for segmenting crowded cellular environments. Using our *in vitro* approach on cellular tomograms, however, did not directly translate. We reasoned that the synthetic data sets were not crowded enough, so we turned to our synthetic cytoplasm again. We trained two single class U-Nets: one network to recognize microtubules and one to recognize ribosomes within a molecularly crowded milieu (Fig. S12). The real neuronal tomogram shown in Fig. 7A was then subjected to segmentation by both networks, as shown in Fig. 7B.

**Figure 7.**
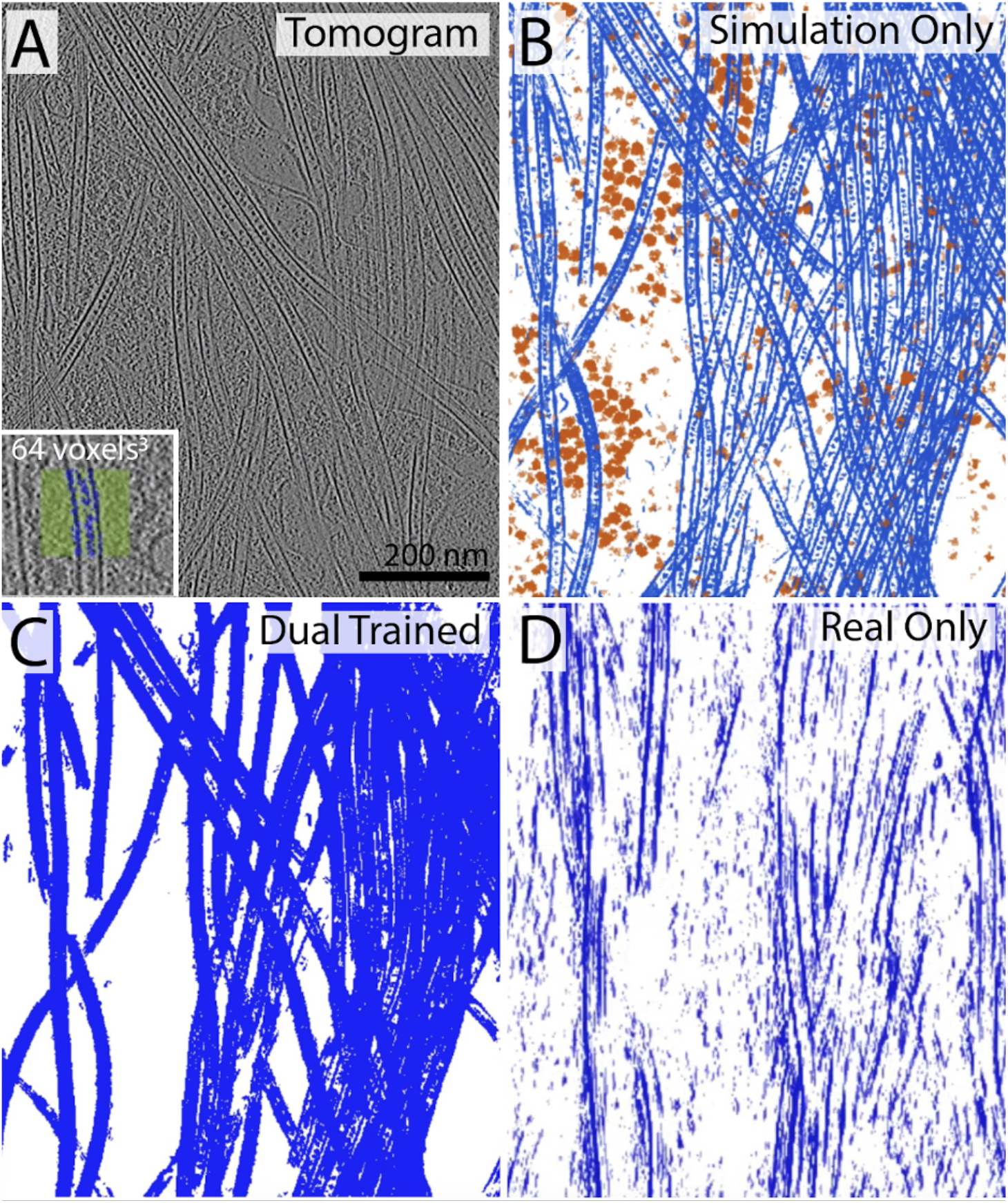
Segmentation of real *in situ* data. A) 2D slice from a neuronal tomographic volume. B) 3D segmentation results for microtubules (blue) and ribosomes (orange) from the tomogram in (A) by two single-class U-nets trained specifically to find each feature within crowded CTS-derived synthetic cytoplasm (shown in Figure S10). C) Segmentation of microtubules with a network simultaneously trained dual trained on the same simulated microtubules plus the 64 cubic voxels of real data shown in the inset of (A). D) Segmentation results from a network only trained on the 64 cubic voxels of real tomographic data

This composite segmentation represents the raw output from both trained networks overlayed and has experienced no postprocessing clean-up.

To test the impact of including hand-annotated real data to the initial training set, we chose a very small amount of data (64×64×64 voxel cube) that included a single stretch of microtubule density (Figure 7A inset) and combined it with our simulated microtubule data to perform what we refer to moving forward as “dual training”. In the dual trained microtubule segmentation, it is clear that the tubule walls are more fully segmented, causing them to lose their transparent appearance as in the “Simulation Only” segmentation. It’s important to note that training only on the small cube of real data did not produce a useful segmentation (Fig. 7D), proving that the addition of even a small amount of real data can fortify networks co-trained with simulated datasets.

We then turned to training multiclass segmentation U-Nets and compared “simulation only” training to our dual trained approach (Fig. 8). For this test we trained a seven-class U-net against our synthetic cytoplasm to recognize actin filaments, cofilactin, microtubules, the chaperonin Tric, ribosomes, synthetic spherical vesicles, and background (see Methods for details). For the dual trained network, the real data used for augmentation was minimal. Each class contributed only a 64 cubic voxel dataset, as depicted in Fig. S13. From the three-dimensional renders (Fig.8A & B), it is simple to conclude that the dual trained network was superior in terms of segmentation noise. Inspection of slices through the segmentations, however, shows that the “simulation only” network performed surprisingly well in recognizing objects from different classes (Fig. 8C & D). While not perfect, we found these results remarkable given that both the data and ground truth could all be generated on a single CPU (from modeling to simulation to reconstruction) in under 30 minutes. Given these results, we currently suggest a dual training strategy for networks meant to segment cellular data, but we believe there is potential for simulations to reach sufficient levels of realism in terms of molecular complexity and noise structure, and this will remain an active area of on-going research.

**Figure 8.**
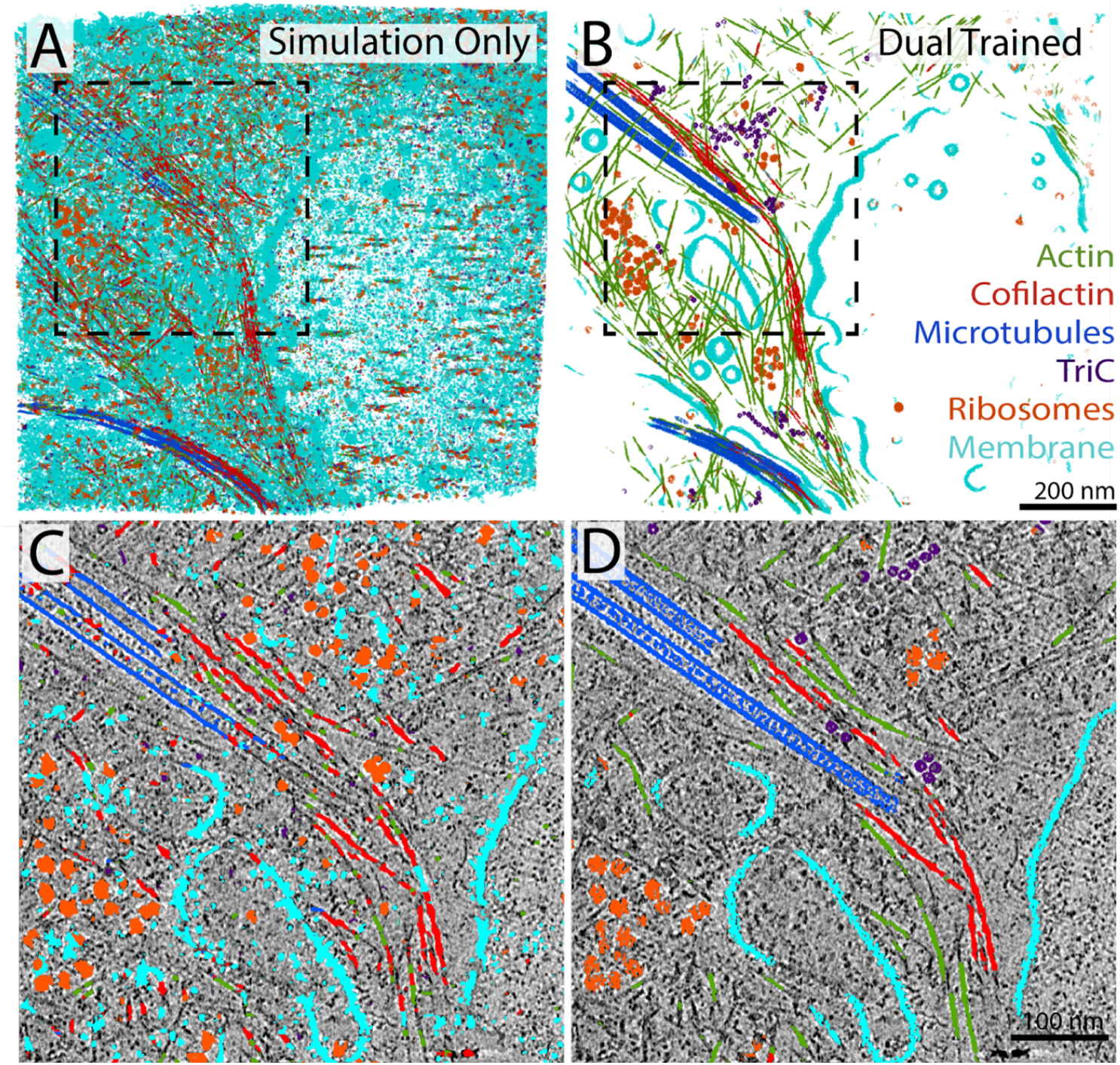
Multi-class segmentation within crowded cellular tomograms. A) 3D segmentation results from a 7-class network (actin, cofilactin, microtubules, TriC, ribosomes, membrane, and background) that was trained solely on these features within crowded synthetic cytoplasm. B) Segmentation of the same tomogram by a U-net dual trained on the same simulated dataset plus six small (64 cubic voxel) hand segmented inputs containing each class (see Fig. S11). C and D) Higher zoom overlays of central a slice through the tomogram with the multi-class segmentation results.

## Discussion

We have showcased CTS, a combined suite for generating models of frozen-hydrated samples and their cryotomographic simulation. It is capable of generating synthetic cryo-ET datasets within minutes from structures of interest, paired with both ground truth segmentations and matched tomograms of ideal to perfect SNR. The simulated datasets and ground truth segmentations can train robust semantic segmentation networks that surpass expert hand segmentations of in-vitro samples, and in cellular samples permit training networks more quickly than manual training, either with retraining or by augmenting with a small amount of target cellular data. Regression denoising is almost impossible to perform without simulated data and has proven to be surprisingly generalizable to different particle mixtures. The intensity of denoising is also manipulable with such easy training, allowing both potent SNR maximizing and gentler denoising that better retains high-resolution information.

Each of these applications highlight how CTS can help fill the gap between the glut of cryo-ET data available and powerful deep learning tools that have the potential to analyze those large datasets. Considering the standard of computer hardware required for collecting and processing cryo-ET data, the technical requirements of CTS are no obstacle, and it can improve resource use by focusing expert microscopists on data collection and analysis rather than manual annotation. Further applications we have not yet explored include training deep learning particle picking tools and directly comparing different reconstruction, denoising, and enhancement techniques quantitatively with known ground truth information. Finally, we have not performed head-to-head comparisons with other deep learning based denoisers or particle classifiers on the same real tomographic data. For instance, it may prove interesting to do in depth comparisons of CTS-trained regression networks to the IsoNet ^21^ approach for missing wedge restoration, or to experiment with the usefulness of training IsoNet networks with simulated data.

While CTS has enabled fast prototyping and powerful deep learning applications, it is deliberately simplified for speed and ease of use. It cannot replace full physics-based all-atom simulators like parakeet ^36^ for quantitative accuracy, especially of very high-resolution data. In the future, CTS could adopt more complete simulation functions to improve rigor and accuracy at the potential cost of longer processing times. More fundamentally, CTS in its current form cannot model ‘true’ cellular or biological superstructures that are often of interest in cryo-ET (mitochondria, Golgi, synapses, etc.), nor does it simulate dynamic molecular properties or interactions.

Despite these limitations, it is remarkable that such small amounts of CTS-generated data can be used to train such powerful and generalizable U-Nets for both image regression and semantic segmentation. Perhaps more important is the ability to derive new training sets, in a matter of minutes, on a single CPU core. This means that anyone can use the wealth of existing structural data (https://www.rcsb.org/, https://alphafold.ebi.ac.uk/, https://opm.phar.umich.edu/, etc.) to quickly iterate different modeling and simulation parameters to systematically test the efficacy of datasets for training neural networks. Additionally, on the modeling side of CTS, users are only limited by their skills and creativity when it comes to building atomic models. Finally, while we have tested the impact of changing basic imaging parameters on a network’s inferability using simulated data, this was only done with U-Net architectures under relatively constrained hyper-parameters. Moving forward, CTS could be used to characterize, optimize, and inform deep learning practices in the cryo-EM field more fully.

## Materials and Methods

### Protein Purification

Purified non-muscle actin (Cat # BK013), cofilin (Cat # CF01-A), and alpha-actinin (Cat # AT01) were purchased from Cytoskeleton, Inc. (Denver, CO). Native Chromatin was isolated and mnase-treated according to the detailed methods described in ^37^. Alpha-CamKII holoenzymes were purified on a CaM-Sepharose column, as described in detail in ^38^.

### Specimen preparation, Image Collection and Reconstruction

Stock actin (10 mg/ml) was diluted 1:10 into ATP-supplemented general actin buffer from Cytoskeleton Inc. (Cat # BSA01-001, 5mM Tris-HCl pH 8.0, 0.2 mM CaCl_2,_ and 0.2 mM ATP), and left on ice for 30 min. Next, 10 ul of actin polymerization buffer (Cat # BSA02-001, 50 mM KCl, 20 mM MgCl_2_, and 1 mM ATP) was added, mixed well, and placed at RT for one hour to polymerize. After polymerization, 2.5 l of Cofilin (5 mg/ml) was added to a tube and diluted in 10 ul of 20 mM Tris-HCl 6.6 Buffer, before adding 12.5 ul of polymerized actin and letting it sit for 30 min at RT. For Alpha-actinin, 40 ul of F-actin was placed in a tube and 10 ul of alpha-actinin (1 mg/ml stock) was added for a final concentration of 200 ug/ml. Add 2 ul of Tris pH 6.5, and leave at RT for 30 min. Before vitrifying CaMKII, stock aliquots dialyzed in 10 mM HEPES, pH 7.4, 0.1 mM EGTA, 200 mM KCl, and 20% glycerol, were thawed and diluted from 3.4 mg/ml to 1 mg/ml in dialysis buffer. Chromatin was prepared according to the detailed protocol in ^37^

All samples were plunge frozen on R2/2 200 mesh Quantifoil EM grids, using a Vitrobot Mark IV (Thermo Fisher). Copper grids were used for purified particles in suspension and gold grids were used for culturing neurons. For purified actin/cofilactin and actin/actinin, chromatin and alpha-CaMKII, 3 microliters of sample were applied to freshly glow-discharged grids and blotted for 3.5 s with force 5. For neurons cultured on grids, we followed the protocol described in detail here ^39^. Briefly, E18 Sprague Dawley rat hippocampi were grown to either DIV 1 or 2 before being picked up in forceps and loaded on the Vitrobot. Grids were hand-blotted through the side port, from the back, using forceps holding blotting paper. After plunge-freezing, samples were stored in liquid nitrogen until they were loaded into the cryo-TEM for tilt series collection.

Tilt series were collected on a Titan Krios (Thermo Fisher) 300 kV cryo-TEM, equipped with a Bioquantum energy filter (Gatan) and either a K2 or K3 direct electron detector. Using Tomography 5 (Thermo Fisher), data was collected from -60 to 60 degrees in either a dose-symmetric scheme or a split scheme with 2-degree increments for everything except purified chromatin, which was imaged with 5-degree tilt increments. Total electron dose was maintained between 60-150 electrons/Å^2^, and defocus was maintained at ∼5 microns. A range of magnifications were used, depending on the sample, from 3.3 - 1.35 Å/pix^2^. Tomographic alignment and reconstruction were carried out using IMOD. All tomograms were reconstructed with weighted back projection and binned 4x before denoising or segmentation.

### PDB structures

See Table S1 for PDBs used and a description of their modifications.

### Generating Synthetic Training Data

#### Modeling Parameters

Tomogram models were generated using the parameters in Table S2. If not listed in the table, the default values were used.

#### Simulation Parameters

Simulations were generated using the parameters in Table S3. If not listed in the table, the default values were used.

#### Ground Truths

For regression networks the ground truth was simulated as tomograms with no CTF corruption or radiation damage, no signal loss at the “detector”, and nearly full angular sampling (−89° to +89° with 1° increments). The full 90 degrees was not used due to limitations imposed by IMOD’s *XYZProject* function. For segmentation networks, the atlases output from CTS during the simulation phase were used as a ground truth for training. The exact specifications of the simulations can be found in Table S3.

### Deep Learning

All U-Net building, training, application, and performance testing were done within the Dragonfly 2022.1 software suite by Object Research Systems. Non-commercial licenses are available through their website at www.theobjects.com, and an open source tutorial video can be found at www.jove.com/t/64435/deep-learning-based-segmentation-of-cryo-electron-tomograms.

In all cases 5-slice U-Nets were used with default architecture and the following hyperparameters: Patch Size = 64 or 128 pixels^2^, Stride Ratio = 1, Batch Size = 32, Epochs = 100-300. The loss function used for regression was Dragonfly’s “MixedORSgradientLoss”, and for semantic segmentation it was “CategoricalCrossEntropy”. For both, the optimization algorithm used was “adadelta”.

### Software Download

CTS can be downloaded from its <u>Github page</u>. The github project also contains installation instructions, required software, and preliminary tutorials on its use.

## Acknowledgements

We’d like to acknowledge Leonardo Ruiz for his assistance in editing this manuscript. This study was supported by the Tobacco Settlement Fund (TSF) grant 4100079742 (to M.T.S), the National Institutes of Health (NIH) grant NS126448-01A1 (to M.T.S), NSF grant MCB-1817929 (to F.A.H.), NSF Grant CHE-220412 (to F.A.H and M.N.W.), NIH Grant R01GM138887 (to F.A.H and M.N.W.), and NSF grant 1911940 (to S.G.). M.N.W. acknowledges the William Wheless III Professorship.

The CryoEM and CryoET Core (RRID:SCR_021178) services and instruments used in this project were funded, in part, by the Pennsylvania State University College of Medicine *via* the Office of the Vice Dean of Research and Graduate Students and the Pennsylvania Department of Health using Tobacco Settlement Funds (CURE). The content is solely the responsibility of the authors and does not necessarily represent the official views of the University or College of Medicine. The Pennsylvania Department of Health specifically disclaims responsibility for any analyses, interpretations, or conclusions.

## References

1. Kreida, S. et al. Cryo-EM structure of the Agrobacterium tumefaciens T4SS-associated T-pilus reveals stoichiometric protein-phospholipid assembly. Structure 31, 385–394.e4 (2023).

2. Swulius, M. T. et al. Structure of the fission yeast actomyosin ring during constriction. Proc. Natl. Acad. Sci. U. S. A. 115, E1455–E1464 (2018).

3. Strauss, M., Hofhaus, G., Schröder, R. R. & Kühlbrandt, W. Dimer ribbons of ATP synthase shape the inner mitochondrial membrane. EMBO J. 27, 1154–1160 (2008).

4. Nicolas, W. J. et al. Cryo-electron tomography of the onion cell wall shows bimodally oriented cellulose fibers and reticulated homogalacturonan networks. Curr. Biol. 32, 2375–2389.e6 (2022).

5. van den Hoek, H. et al. In situ architecture of the ciliary base reveals the stepwise assembly of intraflagellar transport trains. Science 377, 543–548 (2022).

6. Baumeister, W. Cryo-electron tomography: A long journey to the inner space of cells. Cell 185, 2649–2652 (2022).

7. Rodrigues-Oliveira, T. et al. Actin cytoskeleton and complex cell architecture in an Asgard archaeon. Nature 613, 332–339 (2023).

8. Swulius, M. T., Farley, M. M., Bryant, M. A. & Waxham, M. N. Electron cryotomography of postsynaptic densities during development reveals a mechanism of assembly. Neuroscience 212, 19–29 (2012).

9. Gupta, T. K. et al. Structural basis for VIPP1 oligomerization and maintenance of thylakoid membrane integrity. Cell 184, 3643–3659.e23 (2021).

10. Schaffer, M. et al. A cryo-FIB lift-out technique enables molecular-resolution cryo-ET within native Caenorhabditis elegans tissue. Nat. Methods 16, 757–762 (2019).

11. Mageswaran, S. K. et al. In situ ultrastructures of two evolutionarily distant apicomplexan rhoptry secretion systems. Nat. Commun. 12, 4983 (2021).

12. Chreifi, G., Chen, S. & Jensen, G. J. Rapid tilt-series method for cryo-electron tomography: Characterizing stage behavior during FISE acquisition. J. Struct. Biol. 213, 107716 (2021).

13. Eisenstein, F., Danev, R. & Pilhofer, M. Improved applicability and robustness of fast cryo-electron tomography data acquisition. J. Struct. Biol. 208, 107–114 (2019).

14. Peck, A. et al. Montage electron tomography of vitrified specimens. J. Struct. Biol. 214, 107860 (2022).

15. Yang, J. E. et al. Correlative cryogenic montage electron tomography for comprehensive in-situ whole-cell structural studies. bioRxiv 2021.12.31.474669 (2022) doi:10.1101/2021.12.31.474669.

16. Hylton, R. K. & Swulius, M. T. Challenges and triumphs in cryo-electron tomography. iScience 24, 102959 (2021).

17. Xue, L. et al. Visualizing translation dynamics at atomic detail inside a bacterial cell. Nature 610, 205–211 (2022).

18. Hylton, R. K., Heebner, J. E., Grillo, M. A. & Swulius, M. T. Cofilactin filaments regulate filopodial structure and dynamics in neuronal growth cones. Nat. Commun. 13, 2439 (2022).

19. Buchholz, T.-O. et al. Content-aware image restoration for electron microscopy. Methods Cell Biol. 152, 277–289 (2019).

20. Bepler, T., Kelley, K., Noble, A. J. & Berger, B. Topaz-Denoise: general deep denoising models for cryoEM and cryoET. Nat. Commun. 11, 5208 (2020).

21. Liu, Y.-T. et al. Isotropic reconstruction for electron tomography with deep learning. Nat. Commun. 13, 6482 (2022).

22. Chen, M. et al. Convolutional neural networks for automated annotation of cellular cryo-electron tomograms. Nat. Methods 14, 983–985 (2017).

23. Lamm, L. et al. MemBrain: A deep learning-aided pipeline for detection of membrane proteins in Cryo-electron tomograms. Comput. Methods Programs Biomed. 224, 106990 (2022).

24. Moebel, E. et al. Deep learning improves macromolecule identification in 3D cellular cryo-electron tomograms. Nat. Methods 18, 1386–1394 (2021).

25. de Teresa-Trueba, I. et al. Convolutional networks for supervised mining of molecular patterns within cellular context. Nat. Methods 20, 284–294 (2023).

26. Dosovitskiy, A., Fischer, P., Springenberg, J. T., Riedmiller, M. & Brox, T. Discriminative Unsupervised Feature Learning with Exemplar Convolutional Neural Networks. IEEE Trans. Pattern Anal. Mach. Intell. 38, 1734–1747 (2016).

27. Ronneberger, O., Fischer, P. & Brox, T. U-Net: Convolutional Networks for Biomedical Image Segmentation. Med. Image Comput. Comput. Assist. Interv. 234–241 (2015).

28. Wang, F., Wang, H., Wang, H., Li, G. & Situ, G. Learning from simulation: An end-to-end deep-learning approach for computational ghost imaging. Opt. Express 27, 25560–25572 (2019).

29. Kremer, J. R., Mastronarde, D. N. & McIntosh, J. R. Computer visualization of three-dimensional image data using IMOD. J. Struct. Biol. 116, 71–76 (1996).

30. A brief look at imaging and contrast transfer. Ultramicroscopy 46, 145–156 (1992).

31. Santhanam, V., Morariu, V. I. & Davis, L. S. Generalized Deep Image to Image Regression. (2016).

32. Kay, K. The risk of bias in denoising methods: Examples from neuroimaging. PLoS One 17, e0270895 (2022).

33. Swulius, M. T. & Waxham, M. N. Ca(2+)/calmodulin-dependent protein kinases. Cell. Mol. Life Sci. 65, 2637–2657 (2008).

34. Huehn, A. R. et al. Structures of cofilin-induced structural changes reveal local and asymmetric perturbations of actin filaments. Proc. Natl. Acad. Sci. U. S. A. 117, 1478–1484 (2020).

35. McGough, A., Pope, B., Chiu, W. & Weeds, A. Cofilin changes the twist of F-actin: implications for actin filament dynamics and cellular function. J. Cell Biol. 138, 771–781 (1997).

36. Parkhurst, J. M. et al. Parakeet: a digital twin software pipeline to assess the impact of experimental parameters on tomographic reconstructions for cryo-electron tomography. Open Biol. 11, 210160 (2021).

37. Jentink, N., Purnell, C., Kable, B., Swulius, M. & Grigoryev, S. A. Cryo-electron tomography reveals the multiplex anatomy of condensed native chromatin and its unfolding by histone citrullination. bioRxiv 2022.07.11.499179 (2022) doi:10.1101/2022.07.11.499179.

38. Putkey, J. A. & Waxham, M. N. A peptide model for calmodulin trapping by calcium/calmodulin-dependent protein kinase II. J. Biol. Chem. 271, 29619–29623 (1996).

39. Hylton, R. K., Seader, V. H. & Swulius, M. T. Cryo-Electron Tomography and Automatic Segmentation of Cultured Hippocampal Neurons. Methods Mol. Biol. 2215, 25–48 (2021).

