## Supplemental Data for "Rapid Synthesis of Cryo-ET Data for Training Deep Learning Models"

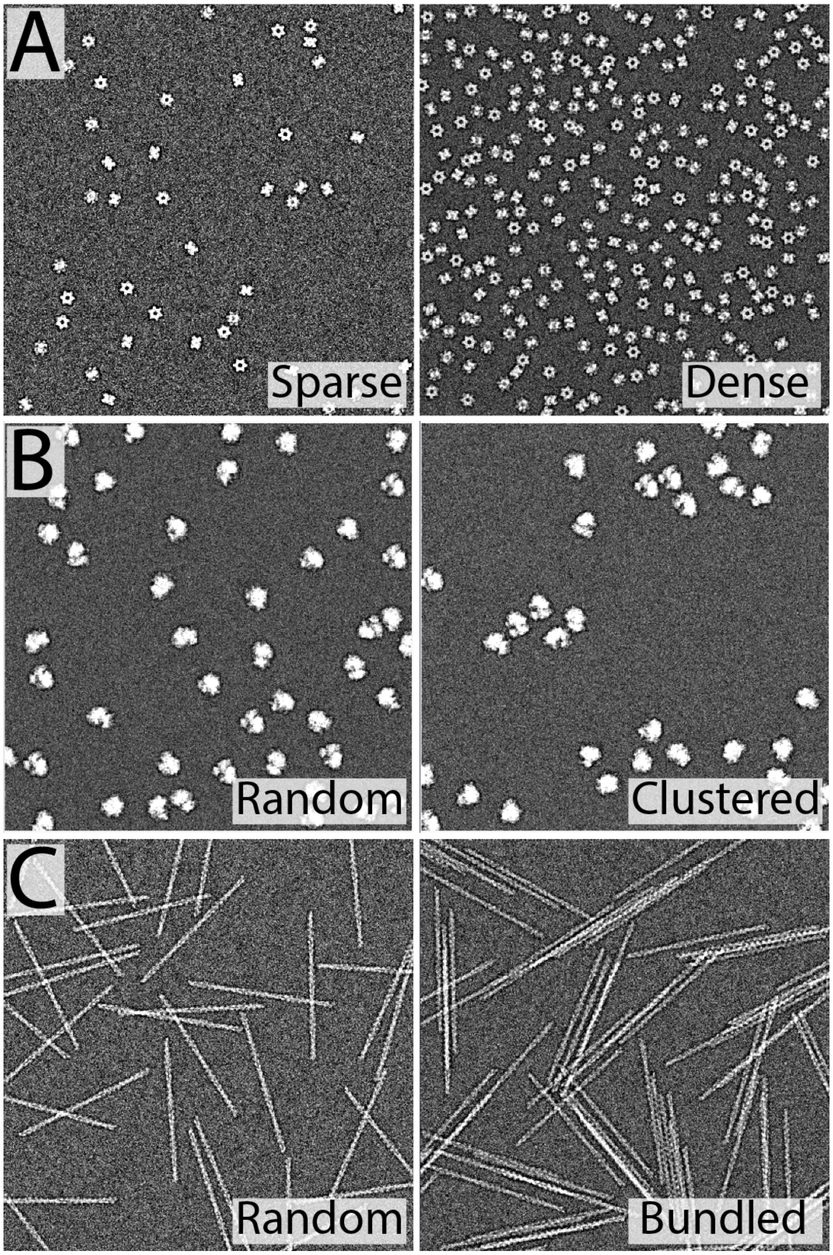


**Figure S1.** Demonstration of some CTS particle placement control. A) Variable density in particle packing of the sample. B) Uniform vs clustered placement of a specific particle within the sample. C) Radial clustering of proteins to enable bundles of thin filaments.

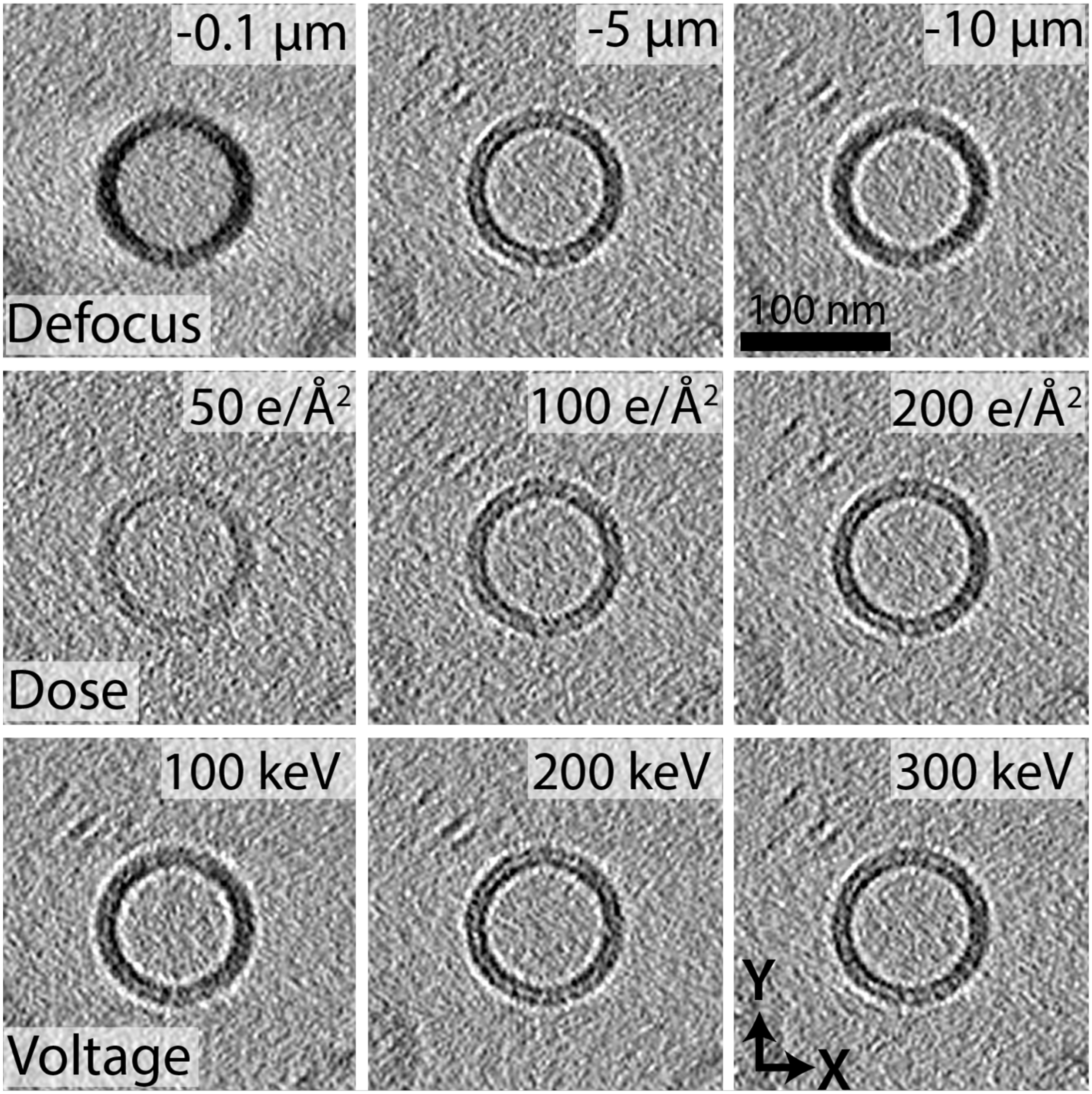


**Figure S2.** Effects of tuning optical parameters in CTS on a simulated tomogram of a modeled spherical vesicle.

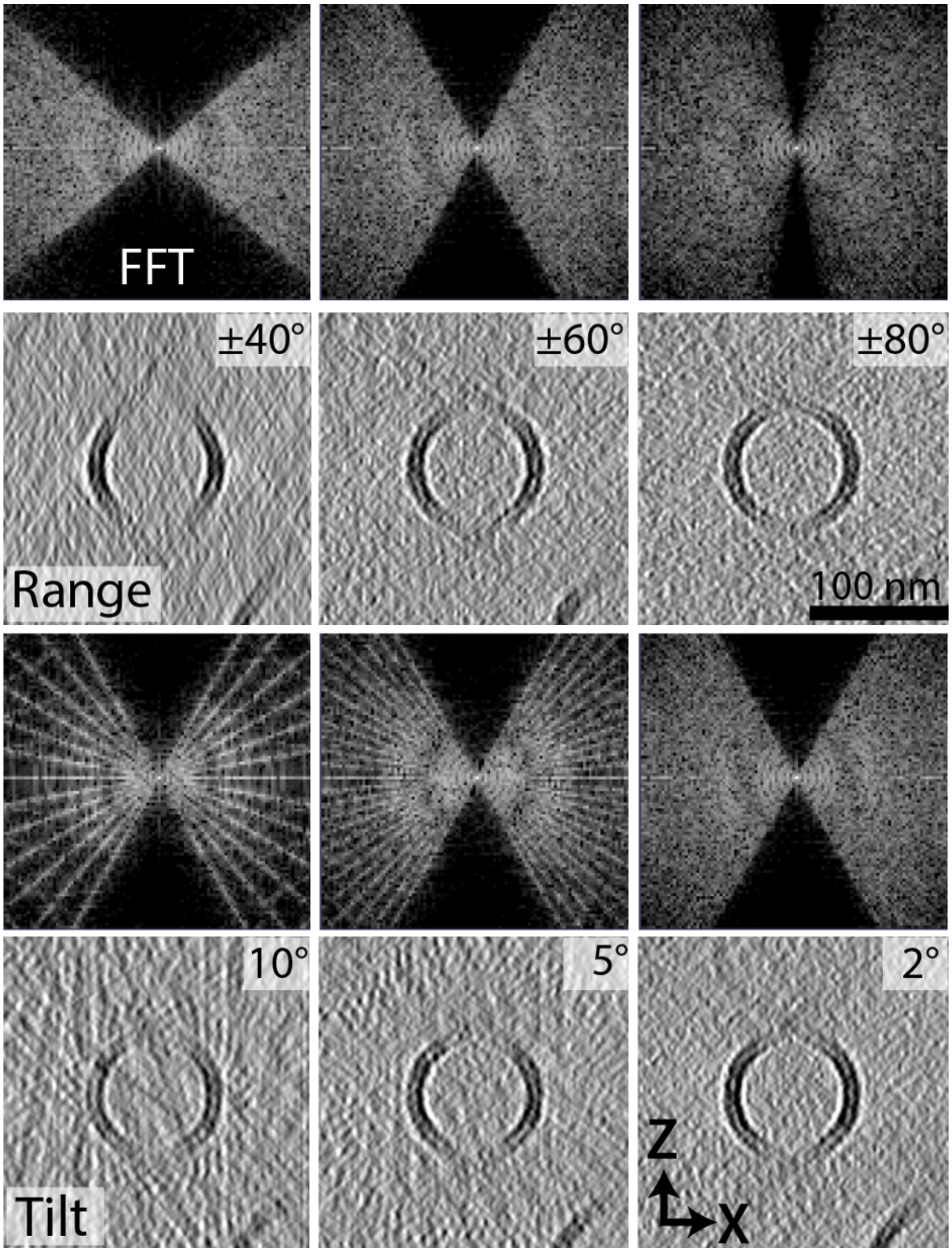


**Figure S3.** Effects of missing wedge range and tilt increment size on a simulated tomogram of a modeled spherical vesicle.

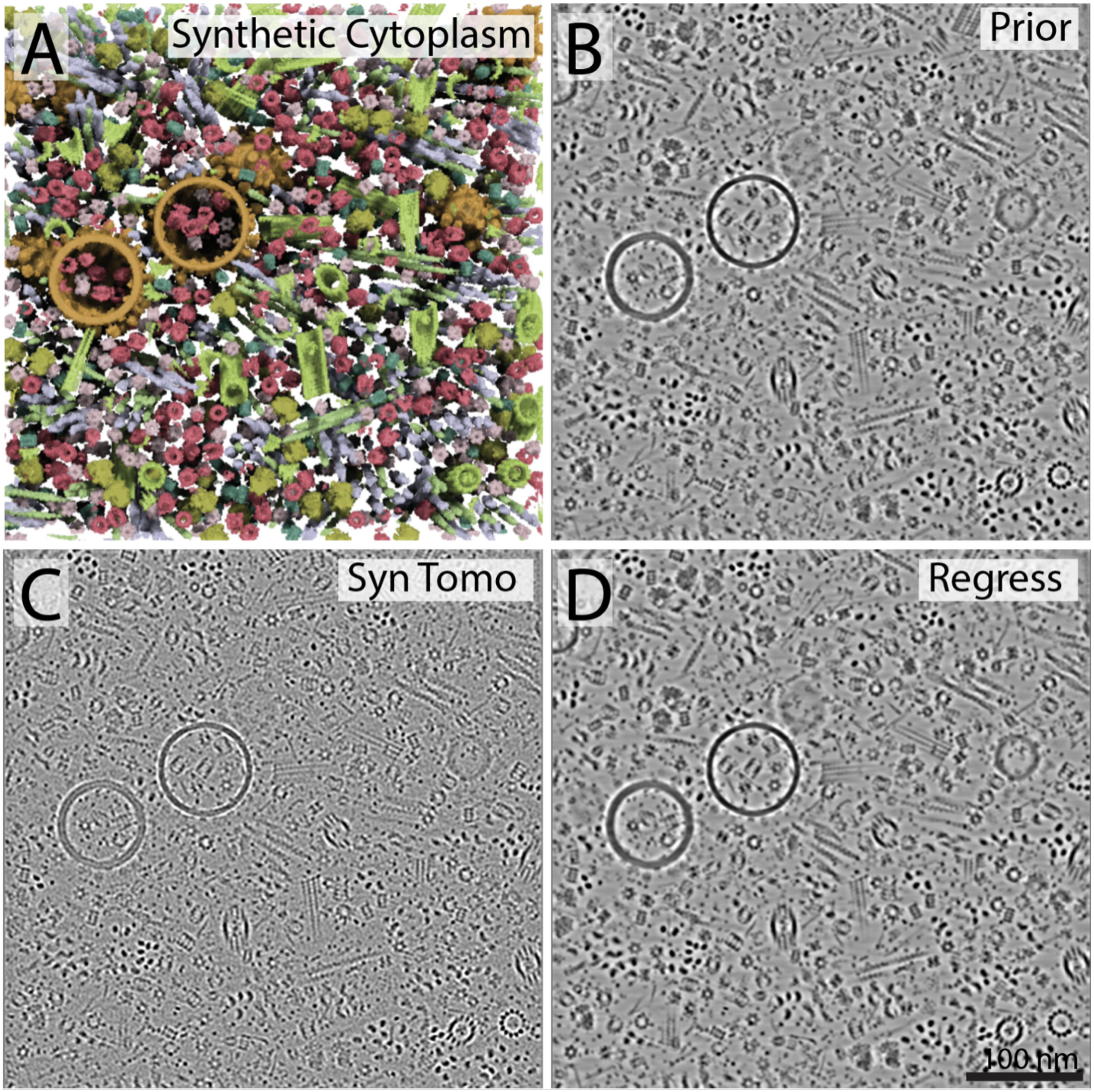


**Figure S4.** Regression of Synthetic Cytoplasm A) Three-dimensional rendering of CTS-generated synthetic cytoplasm used to generate both the tomogram "prior" (B) and the noise/CTF corrupted synthetic tomogram (C). D) U-Net regressed version of the tomogram from panel (C).

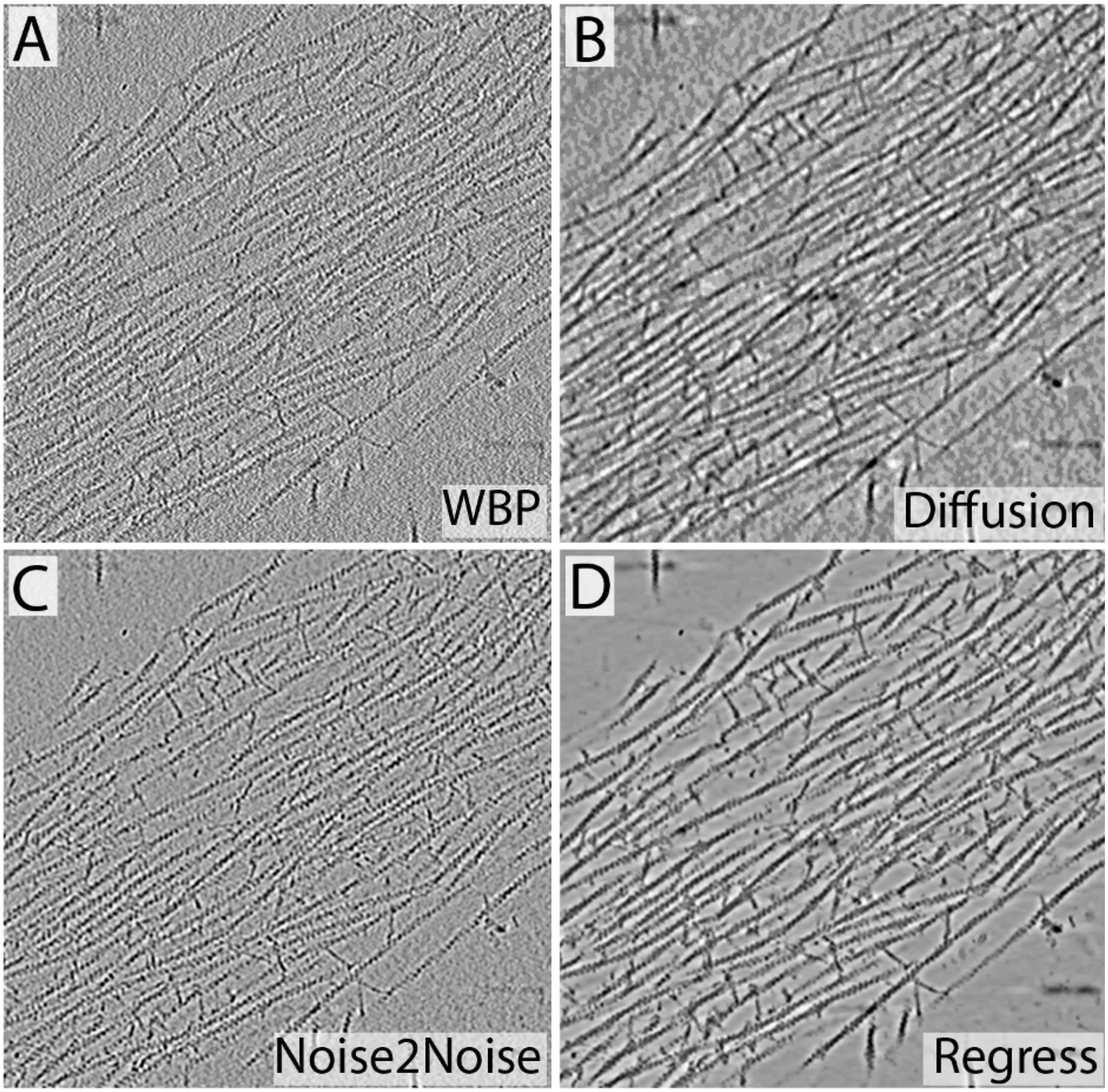


**Figure S5.** Comparison of different denoising strategies. A) Original weighted back projection tomogram. B) Application of anisotropic nonlinear diffusion in IMOD. C) Application of a noise2noise U-Net trained in Dragonfly on two identically simulated tomograms of synthetic cytoplasm with different noise profiles. D) Application of deep learning-based regression with a U-Net trained in Dragonfly on a simulated "prior" with no missing wedge and high SNR.

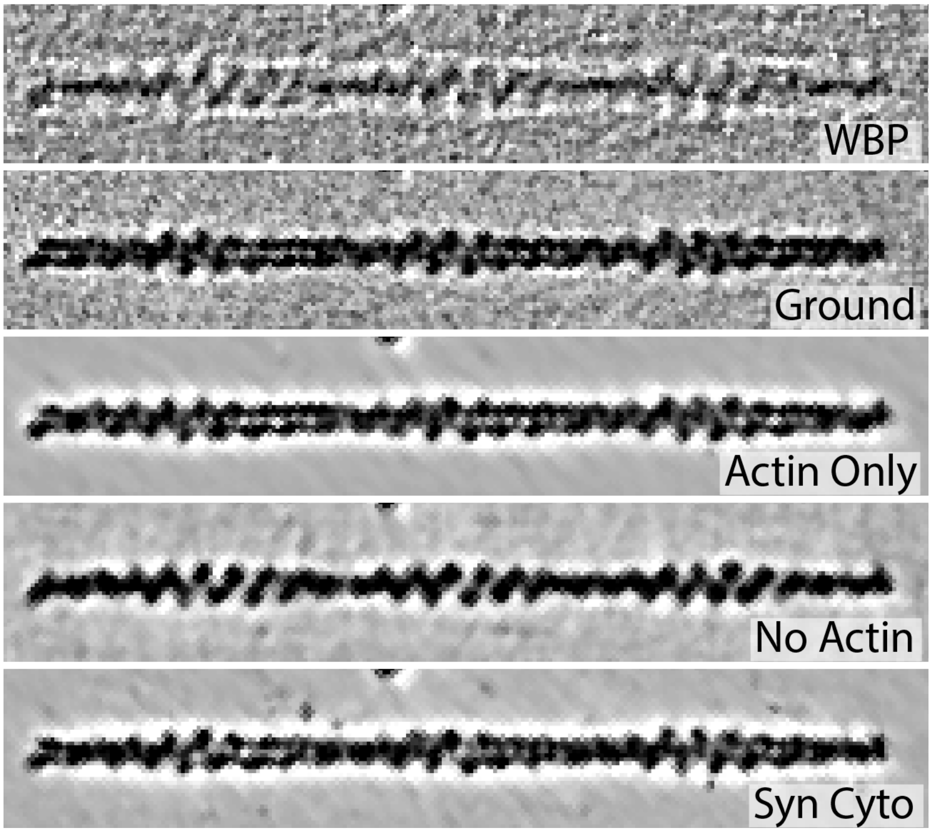


**Figure S6.** The effect of different training inputs on regression of simulated actin filaments. Weighted back projection (WBP) and the high SNR ground truth prior (Ground) shown for comparison. The "Actin Only" network was training solely on actin filaments. The "No Actin" network was trained solely on small globular proteins <200 kDa. The "Syn Cyto" network was trained on synthetic cytoplasm including actin and many other macromolecular complexes (See Methods).

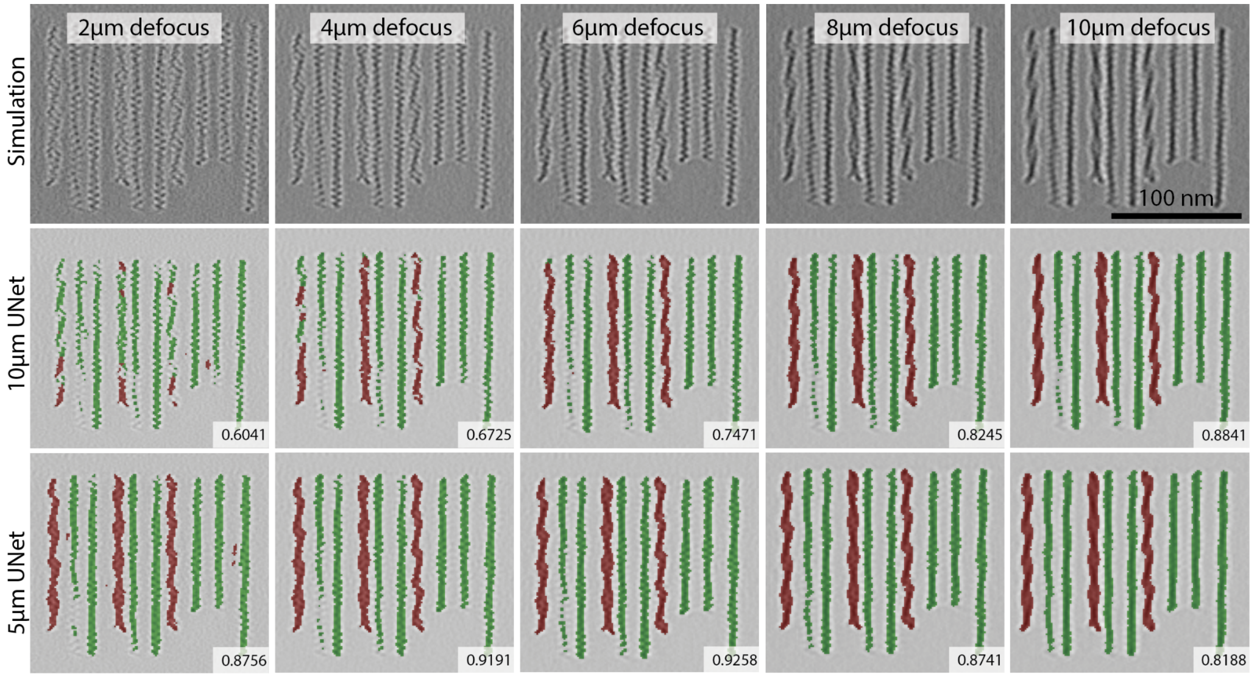


**Figure S7.** Effect of defocus on segmentation accuracy. A simulated tomogram containing rows of actin and cofilactin filaments was generated across a range of defoci from -1 to -10 microns (top row). After training on the -10-micron defocus tomogram the network was inferred to the other nine datasets, where it fell off gradually moving toward -1 micron (middle row). When a network was trained on the -5-micron tomogram it performed well moving in both directions along the gradient (bottom row). Numbers are Dice scores compared to the ground truth.

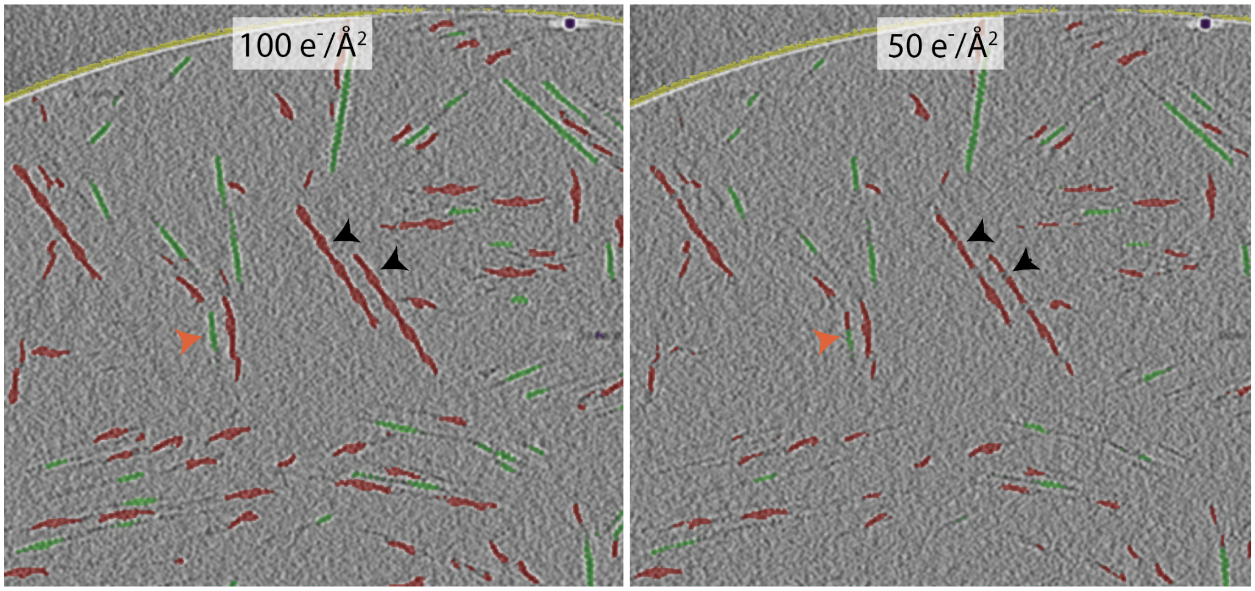


**Figure S8.** Effect of SNR on segmentation accuracy. A U-Net trained on data simulated at 200 $e^-/Å^2$ was inferred to two datasets simulated at lower doses (100 and 50 $e^-/Å^2$). Overall, the inference is robust, but the black arrows highlight regions where the segmentation is noticeably degraded, and the orange arrow highlights where the network has falsely identified a region of transition.

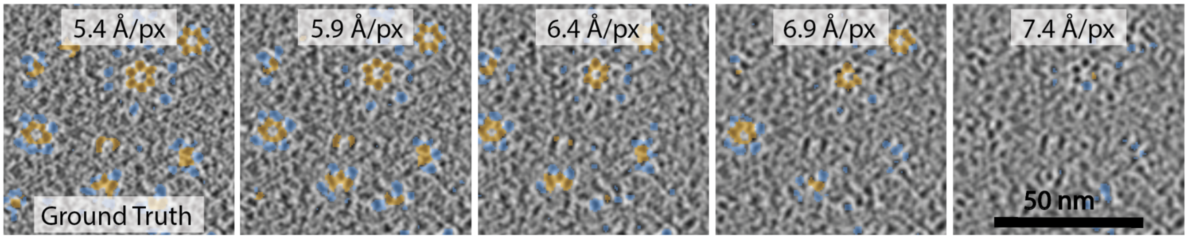


**Figure S9.** Effect of pixel size on segmentation accuracy. A UNet trained to segment the association (orange) and kinase domains (blue) of CaMKII at 5.4 Å/pix loses accuracy when applied to simulations with incrementally coarser pixel sizes. When applied to the same dataset that is simulated at 7.4 Å/pix (far right panel), the model fails completely. The model can tolerate a moderate pixel size discrepancy, but if pushed too far, this parameter significantly affects segmentation quality.

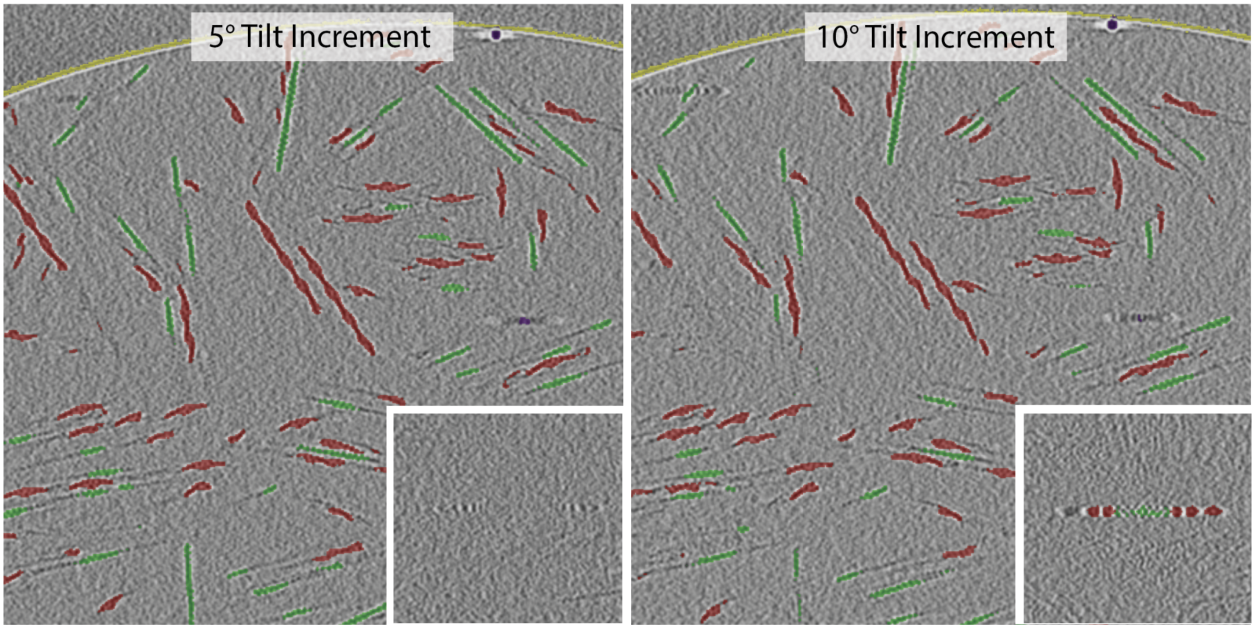


**Figure S10.** Effect of tilt increment on segmentation accuracy. A UNet trained to segment cofilactin (red), and actin (green) with a tomogram simulated at 1° tilt increments infers robustly to tomograms simulated at 5° (left panel) and 10° (right panel) tilt increments. Inset panels demonstrate the primary divergence in accuracy where the artifact signal around a fiducial marker is incorrectly segmented as actin and cofilactin (right inset).

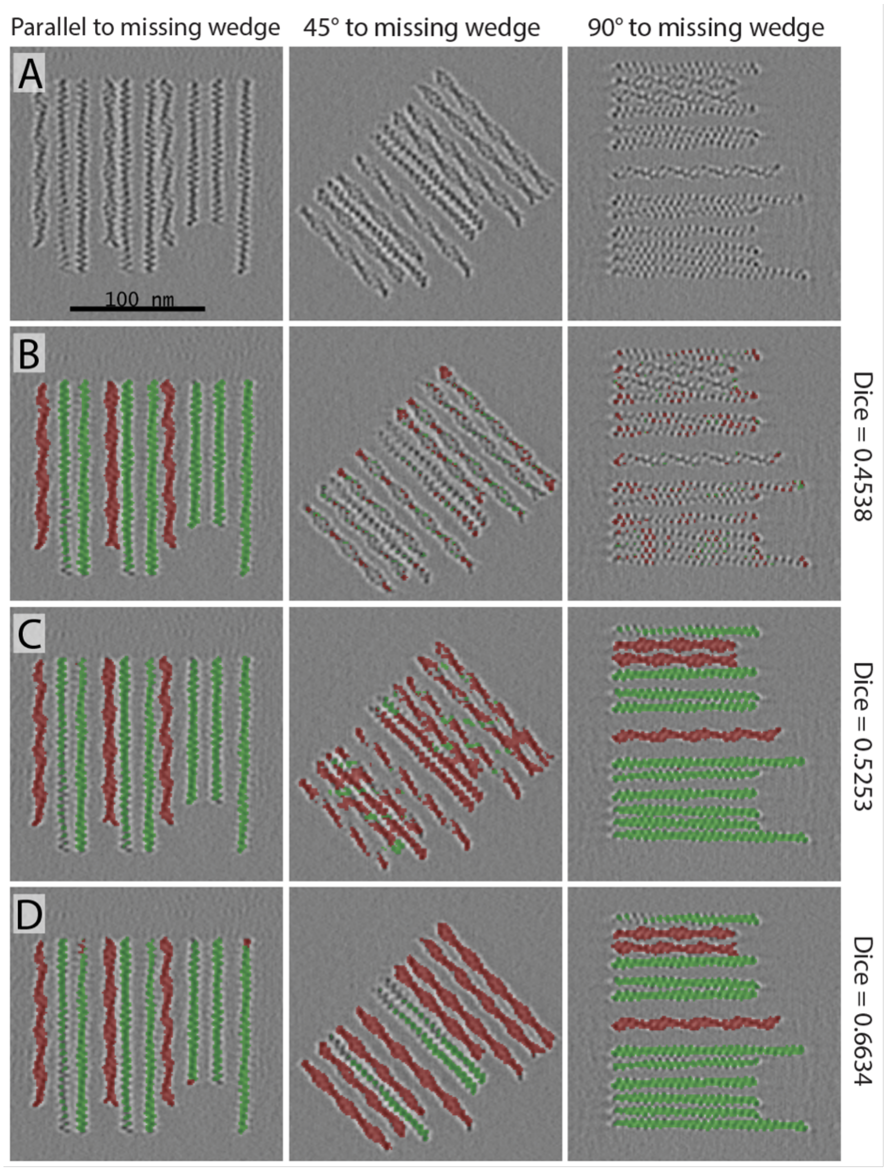


**Figure S11.** Effect of the missing wedge on segmentation accuracy. A) Three panels showing the impact of the missing wedge orientation on the simulated tomogram of actin and cofilactin filaments. B-D) U-Net segmentations of actin (green) and cofilactin (red) by networks trained solely on filaments parallel to the tilt-axis (B), both parallel and perpendicular to the tilt-axis (C), and in all orientations (D).

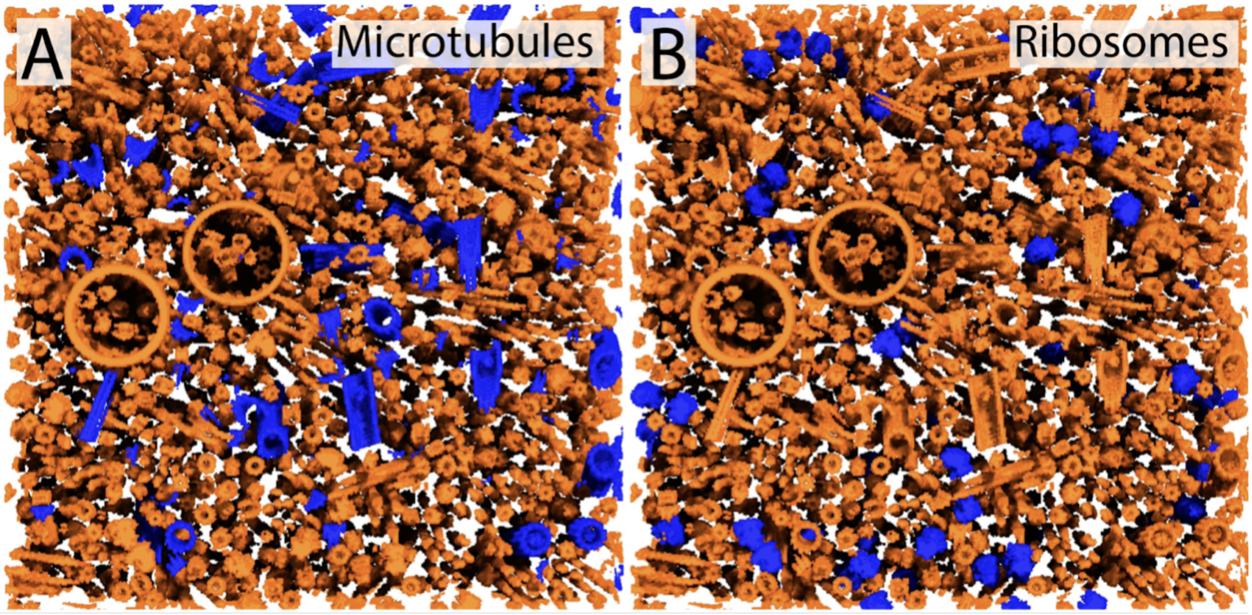


**Figure S12.** Synthetic data used to train two single-class networks to identify and segment microtubules and ribosomes, respectively, in a molecularly crowded milieu.

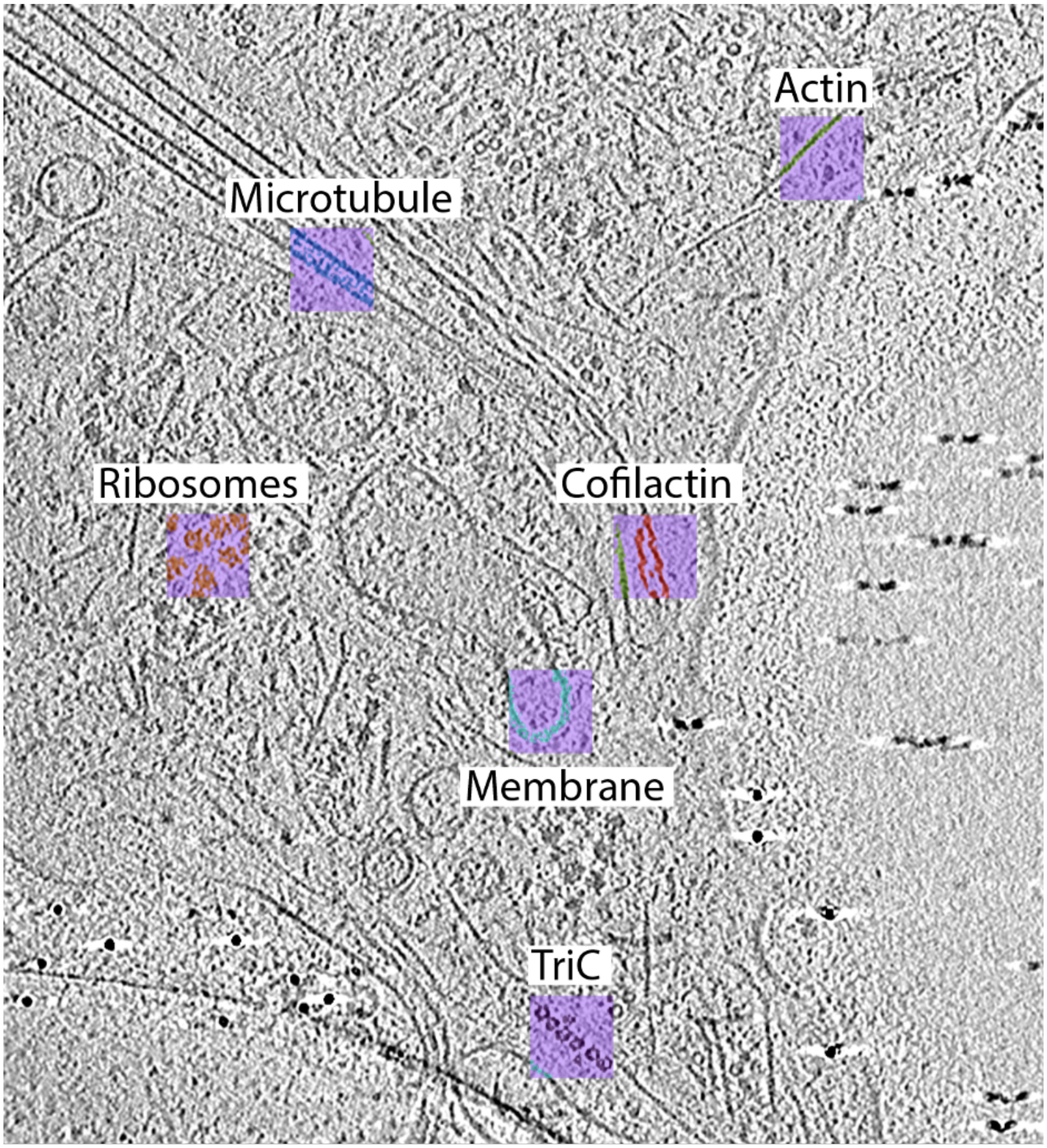


**Figure S13.** Six 64 cubic voxel regions hand segmented and used to augment network training for the results shown in Figure 8B.

| Protein | PDB(s) | Modifications |
|---|---|---|
| Actin Filaments | 6t1y | Elongated helically to 39 subunits |
| Cofilactin Filaments | 3j0s | Elongated helically to 33 subunits |
| Microtubules | 6o2t | Elongated helically by 2x, filled with 3 MIPS |
| Alpha CaMKII Holoenzyme | 3soa, 5u6y | Split into catalytic and association domains |
| NMDAR | 6wht | Split into TM and intra and extra-vesicular domains |
| Adrenomedullin GPCR | 6uva | Split into TM and intra and extra-vesicular domains |
| Proteasome | 5fmg | Split into alpha and beta component submodels |
| TriC | 6ks8 | |
| Ribosome | 4ujd | |
| Distractors | 1qtx, 12gs, 1exr, 1sjj, 2q0u | |
| <200 kDa | 1s3x, 1ul1, 2cg9, 3gl1, 3h84, 3qm1, 4uic, 4wrm, 5a20, 5csa, 5ljo, 6lx3, 6vgr, 6ziu, 7blg, 7blr, 7e6g, 7kfe, 7sk7, 7sgm, 7shk, 7wbt | |
| Chromatin | 1zbb, 5f99, 5oxv, 50y7, 6hkt, 6l4a, 6l49, 6la8, 6m3v, 7pet, 7peu, 7pev, 7pew, 7pex, 7pey, 7pf0, 7pfa, 7pft, 7v9k, 7va4 | |

**Table S1.** PDBs and their modification for simulation.

| Experiment | Figures | Density | Volume | Iterations | Membranes | Pixel Size (Ang) | Molecules |
|---|---|---|---|---|---|---|---|
| Synthetic Cytoplasm Regression | 3, S4 and S5 | 0.8 | 600x600x60 (x3) | 12000 | 5 | 13.2 | Actin Filaments, Cofilactin Filaments, Microtubules, CaMKII, NMDAR, GPCR, Proteasome, TriC, Ribosome, Distractors |
| Chromatin Regression | 4 | 0.4 | 400x400x50 | 1000 | 0 | 8.4 and 6.6 | Chromatin |
| Actin Only Regression | S5 | 0.4 | 400x400x50 | 1000 | 0 | 13.2 | Actin Filaments |
| No Actin regression | S5 | 0.4 | 400x400x50 | 4000 | 0 | 13.2 | <200 kDa |
| Actin and Cofilactin Segmentation | 5 | 0.4 | 400x400x50 | 1000 | 0 | 13.2 | Actin Filaments and Cofilactin Filaments |
| Actin and Cofilin segmentation | 6 | 0.4 | 400x400x50 | 1000 | 0 | 13.2 | Actin Filaments, Cofilactin Filaments modiefied so that Actin and cofilin are separate submodels |
| CaMKII Segmentation | 6 | 0.4 | 400x400x50 | 1000 | 0 | 5.4 | Alpha CaMKII holoenzymes |
| Chromatin Segmentation | 6 | 0.4 | 400x400x50 | 1000 | 0 | 6.6 | Chromatin |
| Proteasome Segmentation | 6 | 0.4 | 400x400x50 | 1000 | 0 | 6.8 | Proteasome |
| In Cellulo Single Class Segmentation | 7 | 0.8 | 600x600x60 | 12000 | 5 | 13.2 | Actin Filaments, Cofilactin Filaments, Microtubules, CaMKII, NMDAR, GPCR, Proteasome, TriC, Ribosome, Distractors |
| In Cellulo Multi Class Segmentation | 8 | 0.8 | 600x600x60 | 12000 | 5 | 13.2 | Actin Filaments, Cofilactin Filaments, Microtubules, CaMKII, NMDAR, GPCR, Proteasome, TriC, Ribosome, Distractors |

**Table S2.** CTS modeling parameters

| Experiment | Figures | Voltage (kV) | Aberration (mm) | Sigma | Defocus (microns) | Tilt | Dose (e/A^2) | Radiation Damage |
|---|---|---|---|---|---|---|---|---|
| Synthetic Cytoplasm Regression | 3, S4 and S5 | 300 | 2.7 | 0.9 | -5 | -60 to +60, 2° | 120 | 1 |
| Chromatin Regression | 4 | 300 | 2.7 | 0.9 | -5 | -60 to +60, 5° | 120 | 1 |
| Actin Only Regression | S5 | 300 | 2.7 | 0.9 | -5 | -60 to +60, 2° | 120 | 1 |
| No Actin regression | S5 | 300 | 2.7 | 0.9 | -5 | -60 to +60, 2° | 120 | 1 |
| Ground Truth Regression | All Regression | 300 | 2.7 | 0.9 | 0 | -89 to +89, 1° | 0 | 0 |
| Actin and Cofilactin Segmentation | 5 | 300 | 2.7 | 0.9 | -5 | -60 to +60, 2° | 120 | 1 |
| Actin and Cofilin segmentation | 6 | 300 | 2.7 | 0.9 | -5 | -60 to +60, 2° | 120 | 1 |
| CaMKII Segmentation | 6 | 300 | 2.7 | 0.9 | -5 | -60 to +60, 2° | 120 | 1 |
| Chromatin Segmentation | 6 | 300 | 2.7 | 0.9 | -5 | -60 to +60, 5° | 120 | 1 |
| Proteasome Segmentation | 6 | 300 | 2.7 | 0.9 | -5 | -60 to +60, 2° | 120 | 1 |
| In Cellulo Single Class Segmentation | 7 | 300 | 2.7 | 0.9 | -5 | -60 to +60, 2° | 120 | 1 |
| In Cellulo Multi Class Segmentation | 8 | 300 | 2.7 | 0.9 | -5 | -60 to +60, 2° | 120 | 1 |

**Table S3.** CTS simulation parameters.
